# A Two-Component Regulatory System Mediates Quorum Sensing–Dependent Morphology and Motility Transitions in the Archaeon *Haloferax volcanii*

**DOI:** 10.1101/2025.11.10.687552

**Authors:** Jacob A. Cote, Priyanka Chatterjee, Marco Garcia, Ran Tao, Arnold Mathijssen, Mechthild Pohlschroder

**Author notes:** Corresponding author: Mechthild Pohlschröder.

## Abstract

Quorum sensing (QS) enables microorganisms—including bacteria, eukaryotes, and viruses—to coordinate collective behaviors in response to population density. Despite their ecological and evolutionary significance, QS mechanisms in Archaea remain poorly characterized. The halophilic archaeon *Haloferax volcanii* provides a model for archaeal QS, transitioning from motile rods to non-motile disks in a density-dependent response to a secreted disk-forming signal (DFS). To identify components of the DFS regulatory network, we screened for spontaneous mutants that retained motility in DFS-containing soft-agar medium. One candidate, *HVO_1357*, encodes a predicted response regulator located adjacent to a histidine kinase (*HVO_1356*) and a second response regulator (*HVO_1358*), consistent with an extended two-component regulatory system (TCS). Based on our results, these genes encode <u>q</u>uorum-sensing <u>a</u>ssociated <u>r</u>egulators (Qar), therefore, we propose rename them *qarA* (*HVO_1357*), *qarB* (*HVO_1356*), and *qarC* (*HVO_1358*). Deletion of *qarA* enabled cells to swim on DFS-containing soft-agar plates and conferred hypermotility on standard soft-agar media; however, these phenotypes were not due to changes in motility-related parameters, but a reduced sensitivity to DFS for induction of the non-motile, disk-shaped state. In contrast, Δ*qarB* and Δ*qarC* strains were non-motile and exhibited premature disk formation during normal growth. Suppressor mutations that restored motility to Δ*qarB* and Δ*qarC* mapped exclusively to *qarA*, suggesting QarA is the central regulator of this system. Transcriptomic analyses revealed that *qarA* deletion leads to upregulation of genes involved in motility and rod-shape formation. Together, these findings reveal *qarABC* as a DFS-responsive regulatory module and represent the first TCS in archaea shown to control QS-dependent behavior.

**IMPORTANCE:** Archaea are ubiquitous and play key roles across diverse ecosystems—including human microbiomes—yet little is known about how they communicate with one another and with other organisms, or how these interactions shape their ecological impact. Such intercellular communication, including quorum sensing (QS), allows microorganisms to coordinate behaviors critical for survival, adaptation, and community organization. In this study, we identify the first archaeal two-component regulatory system that is involved in QS-dependent regulation, providing a foundation for understanding how organisms in this domain sense and respond to population cues. By revealing a previously unknown aspect of archaeal biology, this work represents an important step toward understanding how archaeal communication shapes both their physiology and their interactions within complex microbial communities.

## INTRODUCTION

Microorganisms rely on intercellular communication to coordinate behaviors that promote survival, competition, and adaptation in fluctuating environments. One of the best-characterized mechanisms of microbial communication is quorum sensing (QS), a process that enables density-dependent regulation of gene expression through the production and detection of secreted signaling molecules. When signal concentrations reach a threshold, cells alter transcriptional programs to coordinate behaviors like bioluminescence, biofilm formation, competence, motility, and sporulation (1–4). In gram-negative bacteria, QS often involves acyl-homoserine lactones (AHLs), exemplified by the LuxI/LuxR system in *Vibrio* species, where LuxI synthesizes AHLs that are bound by LuxR, which, once a threshold level is exceeded, activates QS-regulated genes (5). In contrast, gram-positive bacteria typically use secreted peptides as signals, and two-component regulatory systems (TCS) as signal transduction and response pathways (6). QS-like mechanisms have also been identified in fungi and viruses (7, 8), but in archaea quorum sensing remains largely unexplored, representing a major gap in our understanding of microbial communication.

Archaea possess unique physiologies and enzymes with considerable biotechnological potential (9, 10) and, due to their close evolutionary relationship to eukaryotes, serve as powerful models for studying fundamental cellular processes and the origins of eukaryotic biology (11–13). While archaea are recognized as ubiquitous members of environmental microbiomes that play critical roles in global biogeochemical cycles (14–17), they are also increasingly detected in human and animal microbiomes, where their ecological functions and impacts on host health remain poorly understood (18, 19). Despite these remarkable features, archaea remain deeply understudied compared to their bacterial and eukaryotic counterparts. Processes such as quorum sensing and other forms of intercellular signaling may provide mechanisms for communication within archaeal communities in microbiomes, as well as for interdomain communication with bacteria and eukaryotes (20). Understanding the molecular signals and regulatory pathways that mediate these interactions is therefore essential for revealing the broader ecological and physiological roles of archaea across diverse environments.

To date, QS-like behaviors have been reported in only a few archaea. These include *Natrialba magadii*, which exhibits growth phase–dependent Nep protease secretion (21); *Halorubrum lacusprofundi*, where culture supernatants enhance biofilm formation (22, 23); and *Methanosaeta harundinacea*, in which carboxy-AHLs are proposed to mediate density-dependent morphological changes (24). The authors of the latter study also identified a TCS that might be involved in the quorum sensing regulatory pathway, but *in vivo* data have not yet supported this, potentially due to the challenges to genetically manipulate this archaeon (25). In contrast, the model archaeon *Haloferax volcanii* combines a recently identified quorum sensing–like behavior with genetic tractability and facile growth conditions, making it an ideal platform to study archaeal intercellular signaling (20, 26).

In *Hfx. volcanii*, cell morphology changes in a density-dependent manner: low-density cultures are composed of motile, rod-shaped cells, whereas high-density cultures consist of non-motile, pleomorphic disk-shaped cells (27–30). This transition can be artificially induced at low cell densities by adding cell-free conditioned medium (CM), a hallmark of quorum sensing (20). These cell shape transitions are mediated by a secreted, yet-to-be-defined “disk-forming signal” (DFS), which is separable from CM by fractionation and remains bioactive at low concentrations, consistent with small signaling molecule (20, 31). Furthermore, Chatterjee *et al.* identified two *Hfx. volcanii* proteins, DdfA and CirA, as potential components of the DFS regulatory network (20). DdfA (disk-determining factor A) is predicted to function as a shape-associated regulatory protein, as Δ*ddfA* cells remain persistently rod-shaped (32). CirA, a KaiC-like signal transduction protein, shows a similar phenotype, with Δ*cirA* cells also remaining rod-shaped, and has additionally been linked to archaella gene regulation and increased abundance in cells exposed to CM (20, 33, 34). Proteomic data also identified a set of putative TCS proteins that may participate in DFS sensing, analogous to the TCS-based quorum sensing systems seen in gram-positive bacteria (20). One of these proteins is HVO_1356, a sensor histidine kinase encoded near two response regulators: HVO_1357 and HVO_1358. Notably, mutations in *HVO_1357* were previously identified in hypermotile isolates by Collins *et al.* (35), suggesting these proteins may play a role in regulating motility-related responses to DFS.

TCS are among the most versatile and widespread signaling mechanisms in prokaryotes, enabling cells to sense and respond to environmental and intercellular cues through reversible phosphorylation cascades (36–38). Each system comprises a sensor histidine kinase, which autophosphorylates at a conserved histidine residue, and a response regulator that receives the phosphoryl group at a conserved aspartate in the receiver (REC) domain to modulate its activity. The histidine kinase detects various signals—either directly or indirectly—and adjusts the phosphorylation state of the response regulator by balancing its kinase and phosphatase activities. The response regulator in turn orchestrates cellular functions depending on the type of output domain it contains. In *E. coli*, as many as 30 TCS have been experimentally defined (39, 40). In contrast, archaeal TCSs remain poorly understood, with only a few experimental studies reported to date (25, 41–43). Bioinformatic analyses, however, reveal striking differences between bacterial and archaeal TCS components. In bacteria, sensor kinases are typically membrane-bound, response regulators commonly possess DNA-binding domains, and kinases and regulators generally occur in pair. In archaea, by contrast, sensor kinases are often cytosolic, response regulators frequently lack output domains, and kinases commonly outnumber response regulators in the genome (44). These distinctions suggest that archaeal TCSs may have evolved fundamentally different signaling architectures, underscoring the need for further investigation into these systems.

In this study, we used phenotypic screens and reverse genetics approaches to expand our understanding of the DFS regulatory network. Our results show that *HVO_1356*, *HVO_1357*, and *HVO_1358* form a regulatory module that controls motility and cell-shape responses to DFS. HVO_1357 functions as the central regulator, promoting rod-to-disk shape transitions in response to DFS. In contrast, HVO_1356 and HVO_1358 are essential for motility and appear to modulate the activity of HVO_1357. Together, these results identify HVO_1356–HVO_1358 as <u>q</u>uorum-sensing–<u>a</u>ssociated <u>r</u>egulators (Qar) that form a DFS-responsive two-component system mediating quorum-sensing–dependent behaviors in *Hfx. volcanii*—the first such system linked to quorum sensing and among the earliest characterized in Archaea.

## RESULTS

### Identification of mutants less responsive to DFS-dependent motility inhibition

We previously showed that wild-type *Hfx. volcanii* cannot swim on soft-agar plates containing conditioned media (CM), whereas knockout strains of *cirA* and *ddfA* can (20). In an attempt to identify additional proteins involved in sensing or responding to the quorum-sensing signal in CM, we adapted a soft-agar motility assay previously used to identify hypermotile mutants (35), to select for spontaneous mutants that overcome CM-mediated motility inhibition. Mid-log liquid cultures were inoculated as a line across soft-agar plates containing 25% CM. Under these conditions, wild-type cells were non-motile; however, spontaneous mutants with reduced response to DFS regained motility and moved away from the inoculation line, forming flares (Fig. 1). Cells from the edge of such flares were isolated and re-stabbed on CM-containing plates to confirm that they are able to swim in the presence of DFS. Mutants isolated by this procedure were then subjected to whole-genome sequencing. Two independent rounds of this screen yielded 23 isolates, and sequencing identified mutations in 21 of these, including disruptions in *cirA* (five isolates) and *ddfA* (one isolate), validating that our screen can identify genes of interest in the DFS sensing and response pathway (Table 1, full results in Table S1).

**Figure 1.**
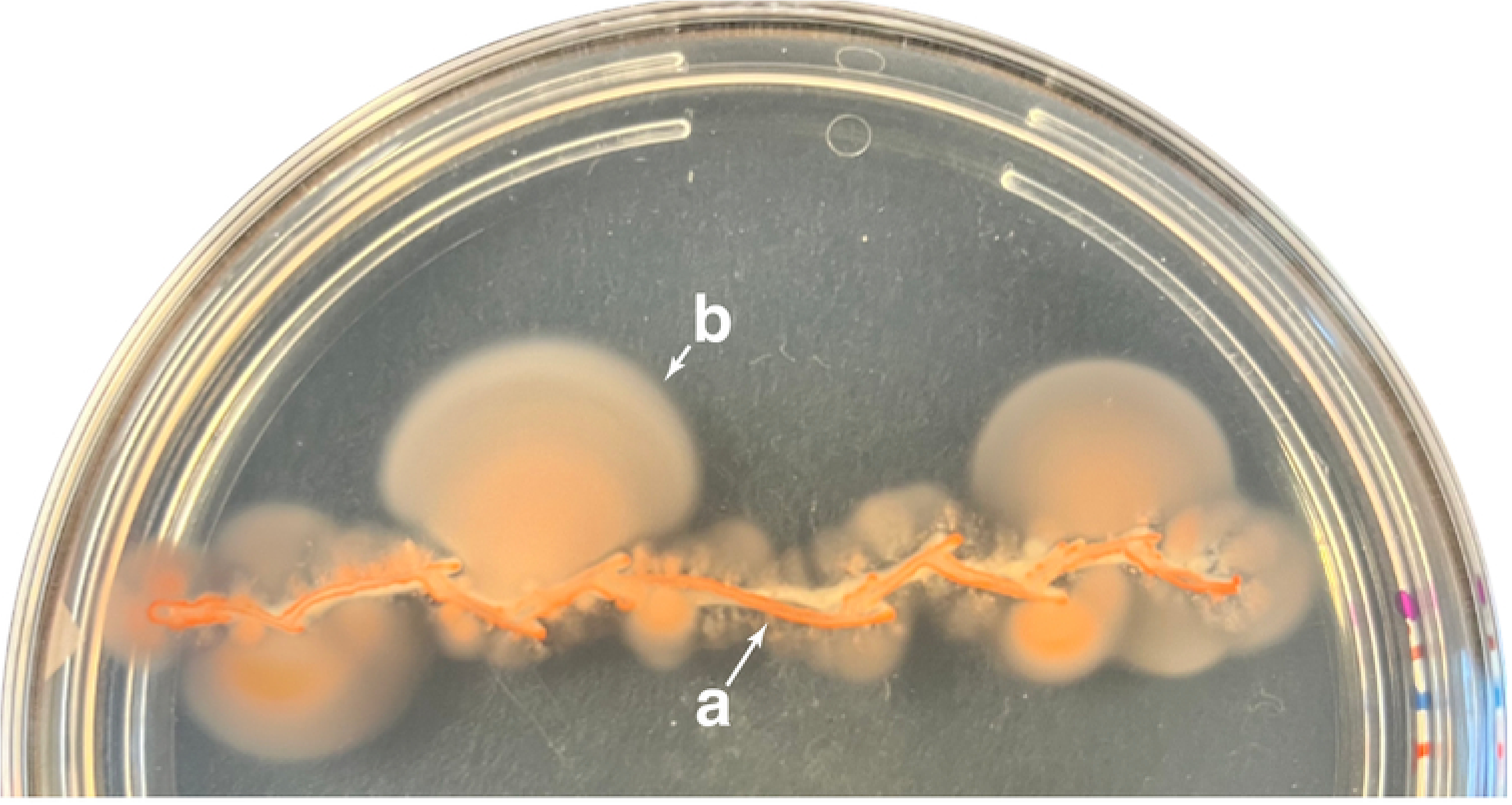
Isolation of mutants that bypass motility inhibition by conditioned medium. Cultures were streaked on 25% CM soft-agar plates and incubated for 4–5 days. For clearer flare observation, this representative plate was stored at room temperature for one week to allow pigmentation of cells to increase. (a) Non-motile wild-type cells (b) Motile flare containing a spontaneous CM bypass mutant.

**Table 1.** Genome alterations in *Hfx. volcanii* mutants less responsive to DFS-mediated motility inhibition.

| Isolate<sup>§</sup> | Affected Gene(s)<sup>†</sup> | Mutation | Position on gene | Secondary Mutation(s)<sup>‡</sup> | Affected gene also identified in *Collins et al. 2018?*<sup>#</sup> | Gene Product |
| --- | --- | --- | --- | --- | --- | --- |
| MG32 | *HVO_1212 (cirA)* | P168L | 505/741 nt | Yes |  | KaiC-type circadian clock protein CirA |
| SB1 | *HVO_1212 (cirA)* | Δ1 bp | 657/741 nt |  |  | KaiC-type circadian clock protein CirA |
| SB7 | *HVO_1212 (cirA)* | +1 bp | 657/741 nt |  |  | KaiC-type circadian clock protein CirA |
| SB13 | *HVO_1212 (cirA)* | F133S | 399/741 nt |  |  | KaiC-type circadian clock protein CirA |
| SB15 | *HVO_1212 (cirA)* | Y43D | 128/741 nt |  |  | KaiC-type circadian clock protein CirA |
| MG6 | *HVO_1357 (qarA)* | L529P | 1586/1749 nt |  | Yes | Receiver/bat box HTH-10 family transcription regulator |
| SB17 | *HVO_1357 (qarA)* | Δ261 bp | 1412-1672/1749 nt |  | Yes | Receiver/bat box HTH-10 family transcription regulator |
| SB18 | *HVO_1357 (qarA)* | +46bp | 1,424/1,749 nt |  | Yes | Receiver/bat box HTH-10 family transcription regulator |
| SB19 | *HVO_1357 (qarA)* | +46bp | 1,424/1,749 nt |  | Yes | Receiver/bat box HTH-10 family transcription regulator |
| MG4 | *HVO_2248* | Δ4 bp | 803-806/816 nt | Yes | Yes | Uncharacterized protein |
| MG7 | *HVO_2239–HVO_2298* | Δ67,424 bp |  |  | Yes | 66 Genes including *HVO_2248* (Uncharacterized protein) |
| SB2 | *HVO_A0350\** | K199N | 597/642 nt | Yes |  | Uncharacterized protein |
| SB20 | *HVO_A0350\** | K199N | 597/642 nt |  |  | Uncharacterized protein |
| MG5 | *HVO_1136 (pgsA1)\** | Δ9 bp | 409-417/825 nt | Yes |  | CDP-alcohol 1-archaetidyltransferase |
| MG8 | *HVO_2176 (ddfA)* | +138bp | 142/201 nt |  | Yes | Disk-determining factor A |
| MG10 | *HVO_2001–HVO_2069* | Δ69,438 bp |  |  |  | 72 genes |
| MG15 | *HVO_2885 (gltP2)\** | A352T | 1,056/1,305 nt |  |  | SDF family transport protein (probable substrate glutamate/aspartate) |
| MG27 | *HVO_2830 (cdc48a)* | D339G | 1,016/2,232 nt |  | Yes<sup>#</sup> | AAA-type ATPase (CDC48 subfamily) |
| SB3 | *HVO_2649* | T→A | -35 nt upstream |  | Yes<sup>#</sup> | Uncharacterized protein |

<sup>§</sup>Full details for each isolate are provided in Table S1.
<sup>†</sup>Mutations predicted by BreSeq and not found in Pohlschröder lab H53 strain, sorted by frequency
<sup>\*</sup>Manual assertion using marginal evidence for isolates where BreSeq did not predict a mutation; see methods for details
<sup>‡</sup>BreSeq-predicted mutations in addition to primary affected gene
<sup>#</sup>Ref. 35; Isolate MC47 contained a Tn-insertion in *HVO\_2649* but had with extensive secondary mutations. MC28, 31 had a Tn-insertion in *HVO\_2248* with an additional T>C (Lys>Glu) mutation within *cdc48a*

One mutant carried a large deletion spanning *HVO_2001–HVO_2069*, which includes genes involved in cell-shape regulation such as *cetZ5*, *volA*, and components of the *agl15*-dependent *N*-glycosylation pathway (32). Several additional mutations were identified in genes previously linked to hypermotility by Collins *et al*. (35), including *cdc48a* (*HVO_2380*, AAA-type ATPase; one isolate), *HVO_2649* (one isolate), and *HVO_2248* (two isolates). This overlap suggests that previously described hypermotility phenotypes may reflect reduced sensing or response to DFS, providing a more specific mechanistic basis for these behaviors beyond general changes in motility regulation. Additional mutants were identified in *HVO_A0350* (two isolates), *pgsA1* (*HVO_1136*, CDP-alcohol 1-architidyltransferase; one isolate), and *gltP2* (*HVO_2885*, SDF family transporter; one isolate). For two isolates, we were unable to identify any mutations despite their ability to swim in the presence of CM, likely due to variant calling challenges associated with polyploid genomes.

In addition to the mutations described above, a final set of mutants highlighted *HVO_1357*, which appeared in four independent isolates; notably, this gene was one of two major loci previously reported by Collins *et al*. to cause hypermotility when disrupted (35). *HVO_1357* is predicted to encode a DNA-binding response regulator and is structurally similar to the Bat protein of *Halobacterium salinarum* (OE3101R), which regulates expression of the bacteriorhodopsin apoprotein (35, 45–47). Unlike Bat, which contains a non-functional REC domain (Fig. S1a) and has yet to be associated with other TCS components, *HVO_1357* appears to function as part of a TCS as it is adjacent to two additional TCS-associated genes: a sensor histidine kinase (*HVO_1356*) and a standalone REC-domain response regulator (*HVO_1358*) (Fig. 2). As demonstrated below, because all three genes in this cluster are related to quorum sensing, we propose naming them <u>q</u>uorum-sensing <u>a</u>ssociated <u>r</u>egulators (Qar), designating HVO_1357—which has the strongest impact—as *qarA*, the histidine kinase HVO_1356 as *qarB*, and the response regulator HVO_1358 as *qarC*.

**Figure 2.**
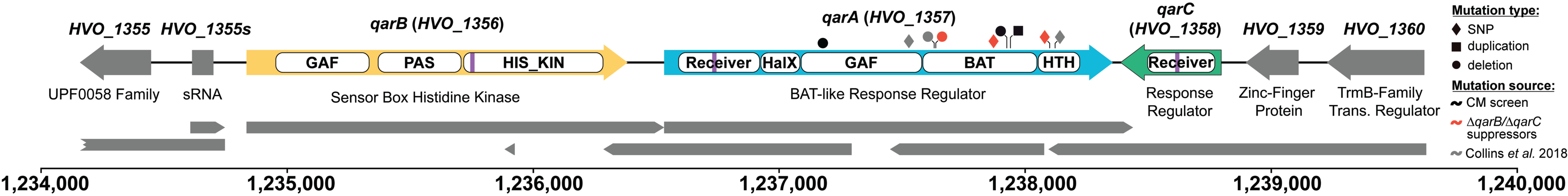
Genomic context, domains, and spontaneous mutations of *qarA*. Arrows representing genes surrounding *qarA* are annotated with locus tag numbers above and protein product names below. Grey arrows indicate the transcript map reported by (54); scale bar and nucleotide numbering are shown below. The types and positions of non-frameshifting *qarA* mutations in this study and by Collins *et al*. (35) are indicated by symbols above *qarA* (key). White boxes within gene arrows denote protein domains identified by InterProScan in the Interpro database (84) for each gene: HVO_1356 (QarB), histidine kinase (D4GXP2); HVO_1357 (QarA), BAT-like response regulator (D4GXP4); HVO_1358 (QarC), response regulator (D4GXP7). Domains include GAF (IPR029016), PAS (IPR035965), HIS_KIN (IPR005467), Receiver (IPR001789), HalX (IPR013971), BAT (IPR031803), and HTH-10 (IPR007050). Purple lines indicate approximate position of phosphoacceptor residues Asp-67 (QarA), Asp-58 (QarC), His-347 (QarB) within their Receiver or HIS_KIN domains respectively.

Multiple sequence alignments with experimentally characterized archaeal and bacterial regulators show QarA contains all key features of a response regulator, including a conserved phosphoacceptor aspartate (D67) and other characteristic motifs of functional REC domains (Fig. S1a)(38). In addition to the REC domain (IPR001789), QarA contains single copies of HalX (IPR013971), GAF (IPR029016), BAT (IPR031803), and HTH_10 (IPR007050) domains, making it a complex multi-domain response regulator (Fig. 2)(35, 48). HalX is haloarchaea-specific and often fused to REC domains (44), while GAF domains typically sense stimuli–such as cyclic nucleotides, small molecules, and light–and appear to be common features of archaeal TCS proteins (44, 49). The BAT domain, of unknown function, is specific to haloarchaea and is typically paired with GAF and HTH_10, a helix–turn–helix DNA-binding domain characteristic of BAT-type regulators (44). Notably, most predicted archaeal DNA-binding response regulators contain BAT domains, and haloarchaeal genomes typically encode multiple BAT-type regulators (44). When gene disruptions in *qarA* from this study and from Collins *et al.* are mapped to these domains, most of the disruptions occur in the BAT domain followed by the GAF and HTH domains (Fig. 2)(35).

### QarA negatively regulates swimming and restricts motility in the presence of CM

To confirm that disruption of QarA induces hypermotility and enables swimming motility in the presence of CM, we constructed a Δ*qarA* strain. No differences in growth were observed by OD₆₀₀ or viable cell counts (Fig. S2a, b). Consistent with phenotypes observed in prior hypermotility screens (35) and our CM-based motility screen, deletion of *qarA* is sufficient to confer hypermotility on soft-agar plates (Fig. 3a) and to permit motility in the presence of CM (Fig. 3b). Complementation of the Δ*qarA* hypermotility phenotype using pTA963 could not be directly assessed, as this commonly used expression plasmid itself induces hypermotility (50, 51). However, in the presence of CM, *qarA* expression *in trans* reduced motility in the Δ*qarA* background compared to the empty vector control, indicating at least partial suppression of the CM-swimming phenotype (Fig. 3d). Notably, pTA963 also enabled wild-type cells to swim in the presence of CM, with motility comparable to pTA963-bearing Δ*qarA* cells, underscoring the strong influence of the plasmid on motility (Fig. 3d). Together, these results suggest that *qarA* expression counteracts the Δ*qarA* phenotype under CM conditions, although full complementation is obscured by plasmid-associated effects.

**Figure 3.**
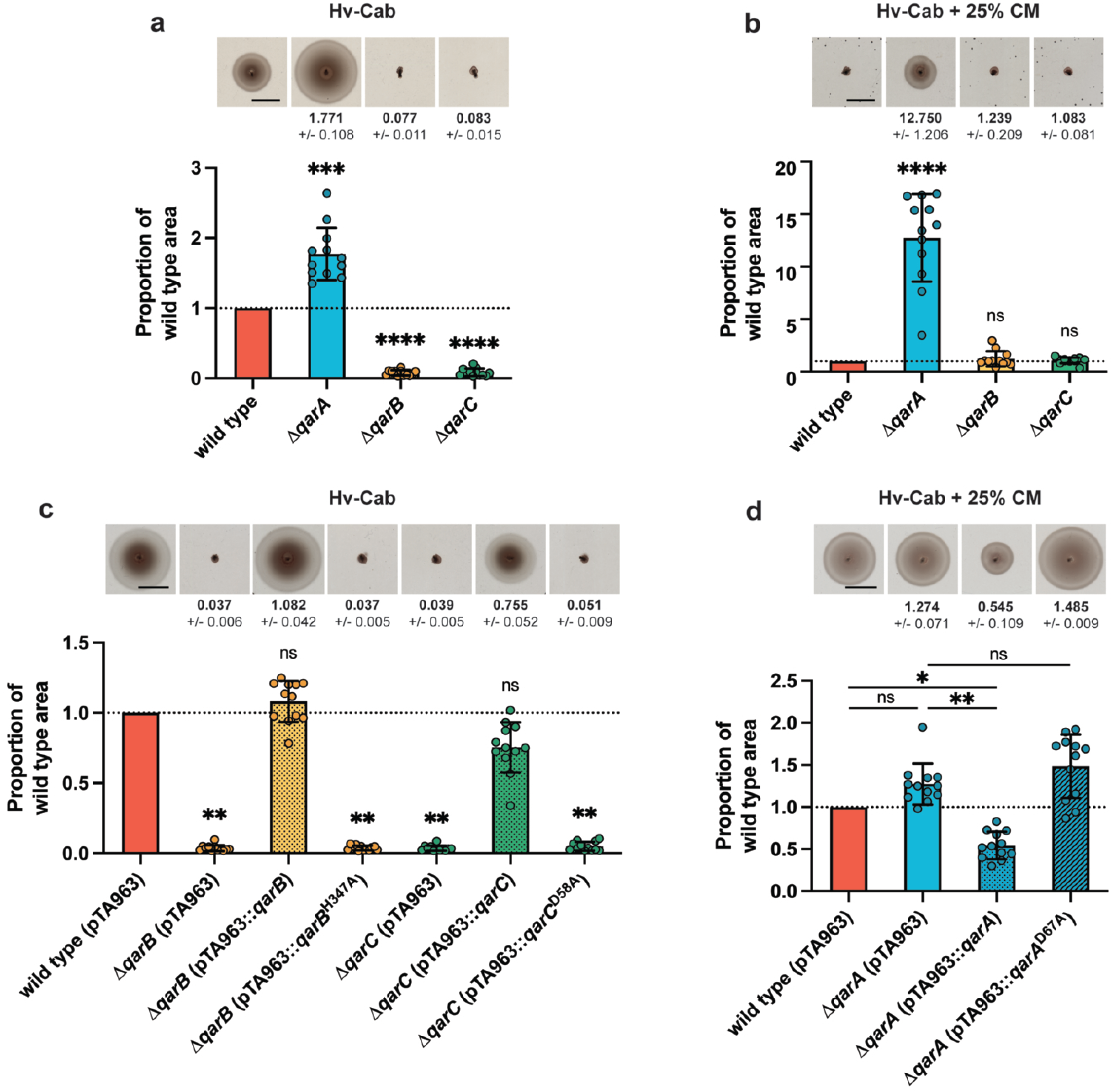
QarABC differentially regulate motility, and key phosphorylation sites are required for their function. Cells were stab-inoculated into 0.35% (w/vol) agar plates containing Hv-Cab medium (a, c) or Hv-Cab medium supplemented with 25% CM (b, d), incubated for 2 days at 45 °C, and then stored at room temperature prior to imaging to allow halo pigmentation to increase. For each plate, halo areas (mm²) were normalized to the wild-type strain on the same plate. Bar graphs show mean values with error bars representing SD and individual data points overlaid; representative halo images of each strain are positioned above their graph and the corresponding numerical mean values (± SEM) are displayed directly below each image. Scale bar, 10 cm. Error bars reflect variation from twelve biological replicates per strain across three independent experiments (24 replicates for wild type in panel c). Statistical analysis was performed on non-normalized halo areas using nested one-way ANOVA, with comparisons to wild type by Dunnett’s test (a–c) or specified pairwise comparisons by Tukey’s test (d). ns, P > 0.05; *P < 0.05; **P < 0.01; ***P < 0.001.

To assess whether loss of *qarA* affects microscopic swimming behaviors that could thereby influence motility on soft-agar plates (52, 53), we used live-cell tracking microscopy to quantify various motility parameters in liquid culture (Movie S1). Consistent with previous studies, we found both wild-type and Δ*qarA* cells displayed two distinct swimming velocities around 2.2*µm*/s and 4.4*µm*/s, corresponding to forward and reverse swimming modes respectively (Fig. 4d)(29). However, we could find no biologically meaningful differences between wild-type and Δ*qarA* cells when comparing other parameters such as average swimming speed (2.197*µm*/s vs 1.975*µm*/s) or motile fractions (48.5% vs 46%) (Fig. 4b, c, d). We also attempted to quantify the average time between directional reversals (e.g. “turns or tumbles”) for both strains; although the standard deviation was high—likely due to background noise—the mean values were similar between wild-type and Δ*qarA* cells and consistent with previously reported values for wild-type *Hfx. volcanii* (Fig. S3)(29). These results suggest that the increased motility of the Δ*qarA* strain is not due to changes in the motility characteristics of any given cell while it is in the motile phase, but rather it may have a defect in regulating when to adopt the motile-rod versus non-motile-disk state.

**Figure 4.**
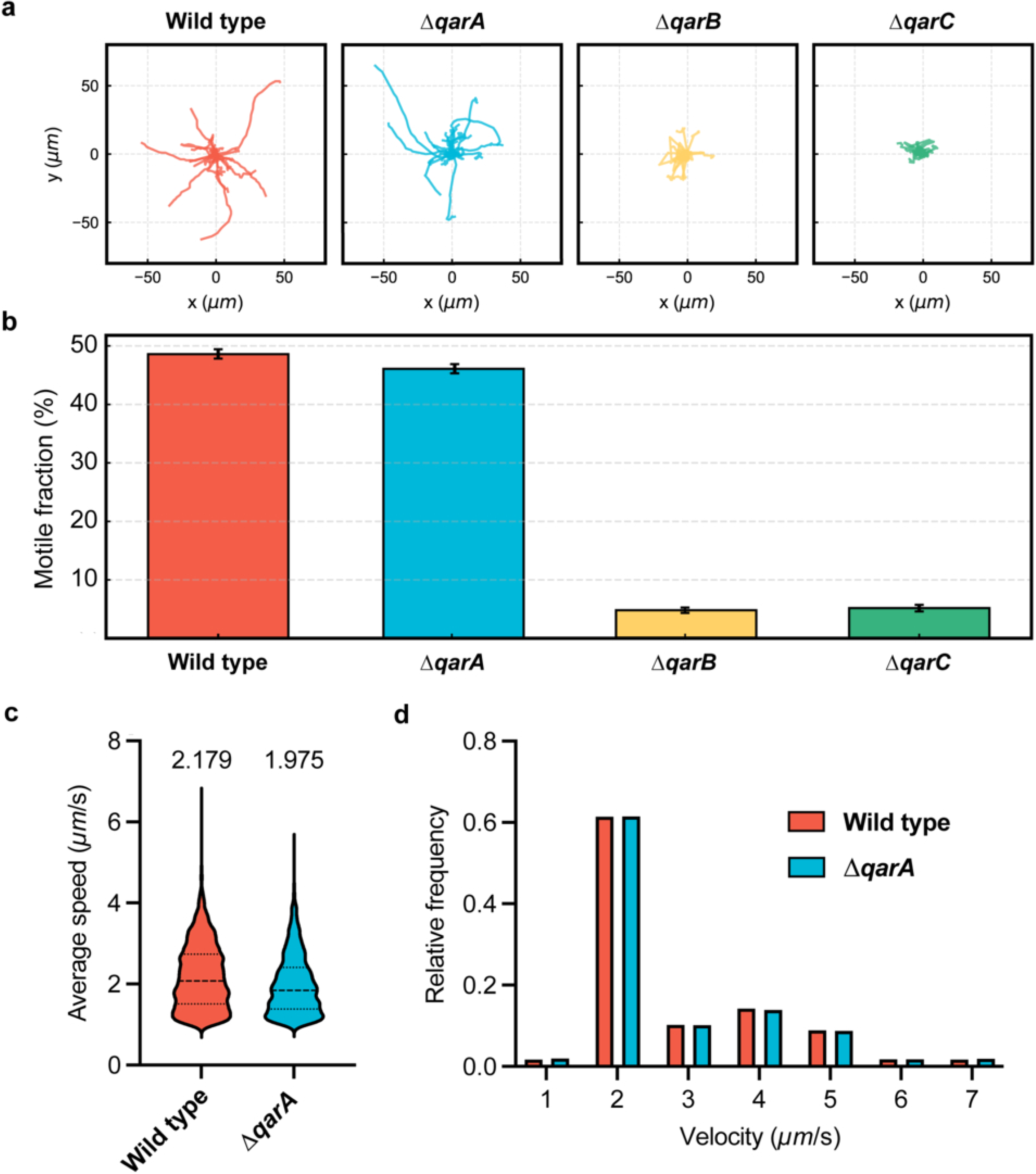
Δ*qarB* and Δ*qarC* are non-motile in early-log liquid culture while Δ*qarA* cells display motility behaviors comparable to wild type. Cells were grown to early-log phase (OD₆₀₀ = 0.045–0.050) and diluted 1:5 (wild type, Δ*qarA*) or 1:2 (Δ*qarB*, Δ*qarC*) in pre-heated motility buffer. Diluted cultures were loaded into pre-prepared motility chambers (see Methods) and imaged immediately under brightfield illumination. Swimming behavior was recorded at 5 fps for 5 min, and trajectories were extracted in FIJI and analyzed using a custom TrackMate-based macro. This resulted in >2000 trajectories for each strain from two independent experiments. (a) Representative single-cell trajectories for each strain. (b) Fraction of motile cells (average swimming speed >1µm/s) within each population, error bars represent SD. (c) Distribution of average swimming speeds for motile strains; average speeds listed above. (d) Relative frequency of instantaneous cell velocities.

### Δ*qarA* raises the DFS threshold for disk formation

We hypothesized that the hypermotility of Δ*qarA* and its ability to swim on CM plates arise from a higher DFS threshold for adopting the non-motile disk shape, resulting in delayed rod-to-disk transition and prolonged motility. To test this, we monitored cell shape across growth phases and following exogenous CM addition. In early-log phase, both wild-type and Δ*qarA* cells were rod-shaped, though Δ*qarA* cells were noticeably longer (average aspect ratios 6.55 vs 5.31)(Fig. 5a, b). By mid-log, many wild-type cells had converted to disks, whereas Δ*qarA* cells remained largely rod-shaped (Fig. 5b). In stationary phase or with 1% (10 µL mL⁻¹) CM, Δ*qarA* cultures fully transitioned to disks, thus precluding a defect in shape transitions (Fig 5b, c, d). We tested our hypothesis by exposing cells to low CM concentrations (0.2–4 µL mL⁻¹) and measuring early-log disk formation. Wild-type cells responded even at 0.2–0.6 µL mL⁻¹ CM, whereas Δ*qarA* required ∼1 µL mL⁻¹ CM to initiate disk formation, reaching wild-type levels by 2 µL mL⁻¹ (Fig. 5e). The shifted response curve demonstrates that deletion of *qarA* raises the initial DFS threshold needed for shape conversion, but does not prevent higher concentrations from triggering the rod-to-disk transition.

**Figure 5.**
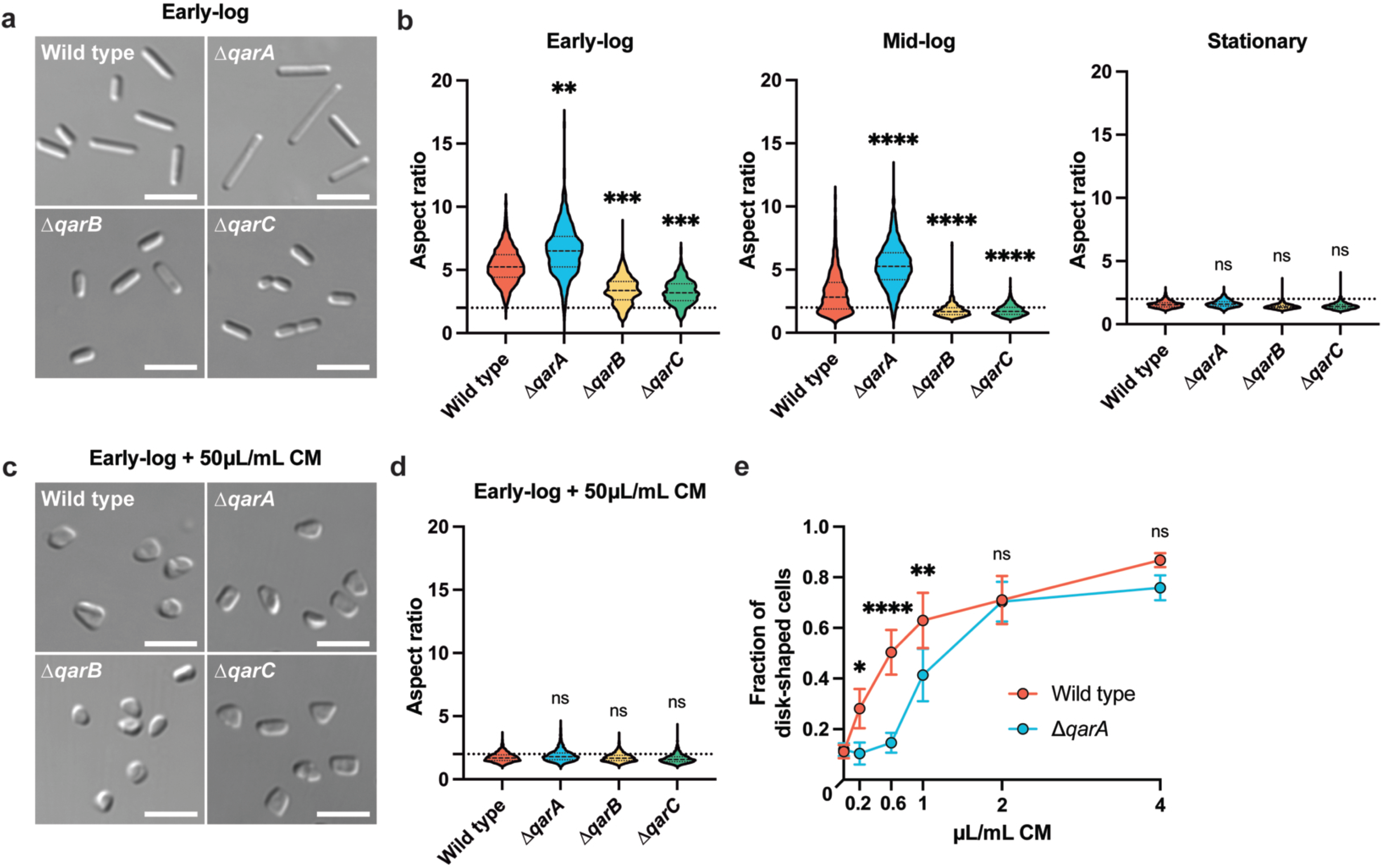
Δ*qarA* cells show reduced sensitivity to DFS signals. Cultures were grown to OD₆₀₀ 0.045–0.050 (early log), 0.30 (mid log), or stationary phase (∼72 h; OD₆₀₀ 1.8–3.0). Differential interference contrast images are shown for (a) early-log cultures without additive and (c) with 1% [vol/vol] CM. Scale bar is 10µm. Cell shape quantification was performed on three biological replicates per strain in three independent experiments for each growth phase without additive (b) or at early-log with 1% [vol/vol] CM (d). Aspect ratio (major axis divided by minor axis) was calculated using CellProfiler; values < 2 (dashed line) indicate disks, and values > 2 indicate rods, with larger ratios reflecting more elongated rods. Statistical analysis performed via nested one-way ANOVA with multiple comparisons to wild type. (e) Wild-type and Δ*qarA* cultures grown to early-log with varying CM concentrations were analyzed for the resulting fraction of disk-shaped cells using CellProfiler eccentricity values (foci of ellipse-shaped object divided by major axis); values <0.9 indicate disks, and >0.9 indicate rods. Error bars represent SD of three biological replicates per strain across three independent experiments. Statistical analysis performed via two-way ANOVA with Šídák’s multiple comparisons test. ns P > 0.05; *P < 0.05; **P < 0.01; ***P < 0.001; ****P < 0.0001.

### QarB and QarC are required for motility and maintenance of the motile rod shape

Previous bioinformatic analyses revealed that *qarB*, *qarA*, and *qarC* are located in the same genomic region but are not organized as an operon (35, 54, 55). *qarB* encodes a putative sensor histidine kinase with conserved PAS and GAF sensory domains but no transmembrane regions, suggesting it functions as a cytosolic sensor or interacts with other membrane-bound components for signal transduction through the cell membrane (35, 48). *qarC* encodes a standalone response regulator containing a conserved receiver domain but lacks an output domain—an arrangement more frequently observed in archaea than bacteria (44, 48). Both *qarB* and *qarC* encode the essential phosphoacceptor residues H347 and D58 respectively, as well as other residues important for histidine kinase and REC domain functions (Fig. S1a, b)

To define their roles in motility and cell shape, deletion mutants were generated and compared to wild type and the Δ*qarA* strain. While Δ*qarB* and Δ*qarC* showed slightly lower OD₆₀₀ values, the viable counts at comparable growth times remained similar (Fig. S2a, b). However, motility phenotypes of these mutants were striking: both were completely non-motile on soft agar with or without 25% CM (Fig. 3a, b). Complementation with plasmid-borne *qarB* or *qarC* restored motility to wild-type levels, confirming that both genes are essential for motility (Fig. 3c).

Microscopy imaging revealed that in early log phase, Δ*qarB* and Δ*qarC* cells were rod-shaped, yet significantly shorter than wild type (average aspect ratios 3.50 and 3.32 vs 5.31) and transitioned to disk morphology earlier in growth (Fig. 5a, b). Although still rod-shaped, live-cell microscopy showed that these rods were non-motile under conditions where wild-type and Δ*qarA* strains were motile (Fig. 4a, b). When grown in the presence of CM, Δ*qarB* and Δ*qarC* cultures were fully disk-shaped (Fig. 5c, d). Together, these results show that *qarB* and *qarC* are both required for motility and may promote the motile cell state.

### QarABC exhibits features of a TCS

Given the domain architectures and sequence features consistent with TCS proteins, we next asked whether QarABC functions as a canonical phosphotransfer-based signaling system. Specifically, we tested whether the conserved phosphoacceptor residues required for response regulator and histidine kinase activity are necessary for QarABC function. The phosphoablative mutant *qarA*^D67A^ expressed from pTA963 failed to suppress the CM-swimming phenotype of the Δ*qarA* strain, indicating that QarA functions as a response regulator (Fig. 3d). Similarly, phosphoablative mutants QarB^H347A^ and QarC^D58A^ also abolished complementation (Fig. 3c), suggesting that all three proteins function as components of a TCS-like signaling pathway.

Consistent with a functional relationship among QarABC, spontaneous motile suppressors emerged from both Δ*qarB* and Δ*qarC* strains (Fig. 4a). Whole-genome sequencing of three suppressors from each background solely revealed mutations in *qarA* (Table S1), indicating that loss of *qarA* restores motility in these non-motile mutants and functionally links QarA to QarB and QarC.

Motivated by these observations, we performed epistasis analysis to further define the QarABC pathway architecture. We generated a Δ*qarBC* double mutant and used suppressor strains in which Δ*qarB* and Δ*qarC* backgrounds carry mutations in *qarA* as functional proxies for Δ*qarAB* and Δ*qarAC* double mutants. Disruption of *qarA* suppressed the non-motile phenotypes associated with loss of either *qarB* or *qarC*, indicating that *qarA* is epistatic to both genes (Fig. 6b, 3a). Moreover, the Δ*qarBC* double mutant displayed a non-motile phenotype comparable to the single Δ*qarB* and Δ*qarC* mutants, consistent with QarB and QarC acting within the same pathway (Fig. 6b, 3a).

**Fig. 6.**
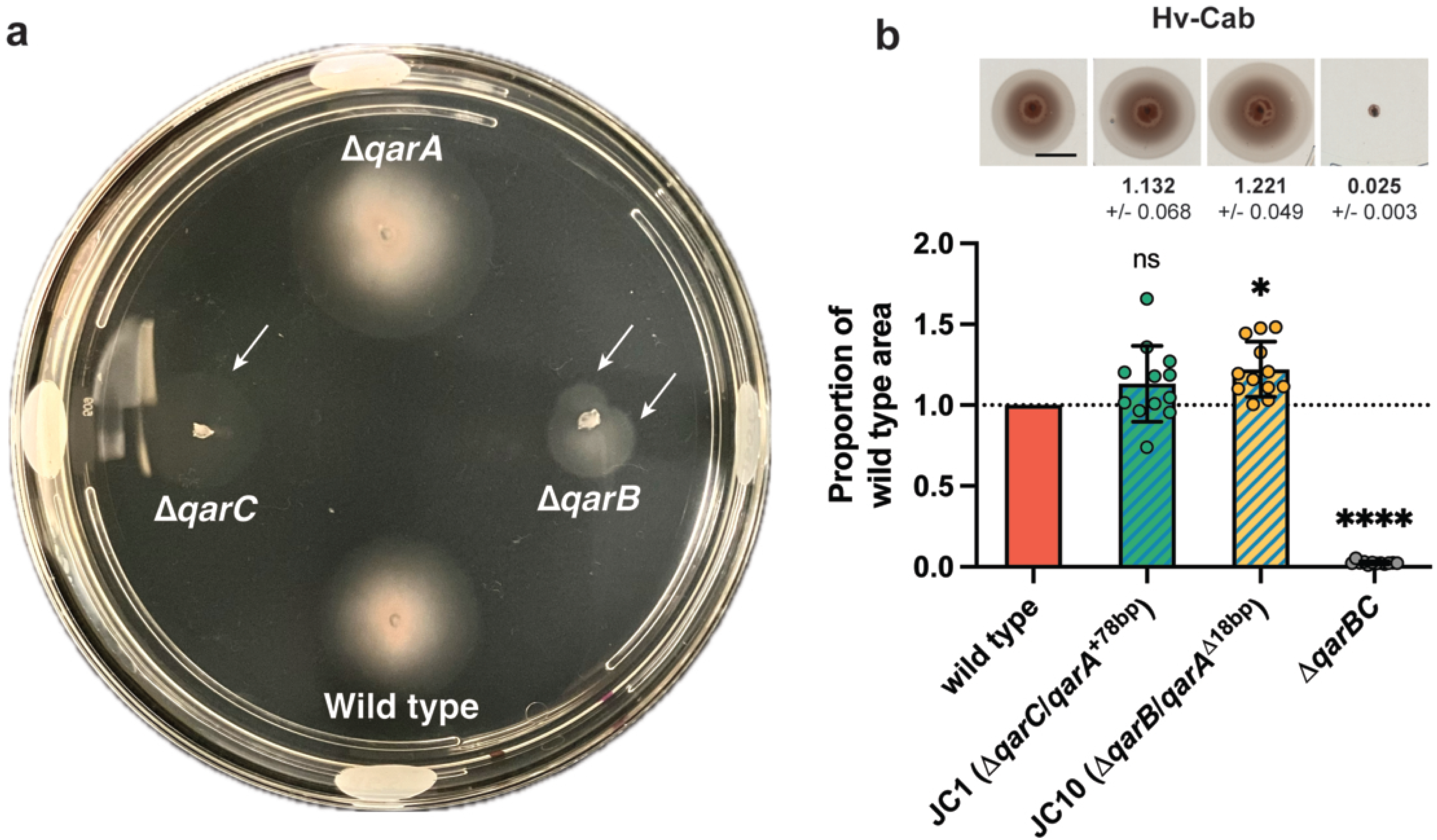
QarA is epistatic to QarB and QarC, which act in the same pathway. Representative soft-agar plate displaying motile suppressors of Δ*qarB* and Δ*qarC*, with white arrows indicating motility halos formed by spontaneous suppressor mutants (a). Epistasis analysis of *qarABC* genes using a Δ*qarBC* deletion strain and Δ*qarB* or Δ*qarC* spontaneous suppressor mutants which contain mutations in *qarA* (b). Cells were stab-inoculated into 0.35% (w/vol) agar plates containing Hv-Cab medium, incubated for 2 days at 45 °C, and then stored at room temperature prior to imaging to allow pigmentation to increase. For each plate in the epistasis assay, halo areas (mm²) were normalized to the wild-type strain on the same plate. Bar graphs show mean values with error bars representing SD and individual data points overlaid; representative halo images of each strain are positioned above their graph and the corresponding numerical mean values (± SEM) are displayed directly below each image. Scale bar, 10 cm. Error bars reflect variation from twelve biological replicates per strain across three independent experiments. Statistical analysis was performed on non-normalized halo areas using nested one-way ANOVA, with comparisons to wild type by Dunnett’s test. ns, P > 0.05; *P < 0.05; ****P < 0.0001.

### *qarA* influences transcription of motility, cell-shape, and TCS genes

To identify genes differentially transcribed in the absence of the putative DNA-binding response regulator QarA, we compared RNA-seq profiles of wild-type and Δ*qarA Hfx. volcanii* at early-log and late-log phases (full datasets in Tables S2–S5).

#### Early-log phase

In Δ*qarA*, 60 differentially expressed genes (DEGs) were identified (Fig. 7a; Table S3). Thirteen DEGs were upregulated, including genes encoding archaella components *arlA1* (*HVO_1210*) and *arlA2* (*HVO_1211*), motor complex components *arlH* (*HVO_1216*) and *arlI* (*HVO_1217*), and the chemotaxis adaptor *cheF* (*HVO_1219*), all of which are important for motility (33, 56). Genes encoding proteins involved in rod formation, *rdfA* (*HVO_2174*) and *sph3* (*HVO_2175*), were also upregulated (32). *HVO_2649*, identified in our 25% CM screen as well as the previous hypermotility screen (35), was similarly upregulated. The most highly upregulated genes were *HVO_B0192* and *HVO_B0193*, encoding a putative transmembrane protein and transcriptional regulator with a domain of unknown function (DUF7521), respectively.

**Figure 7.**
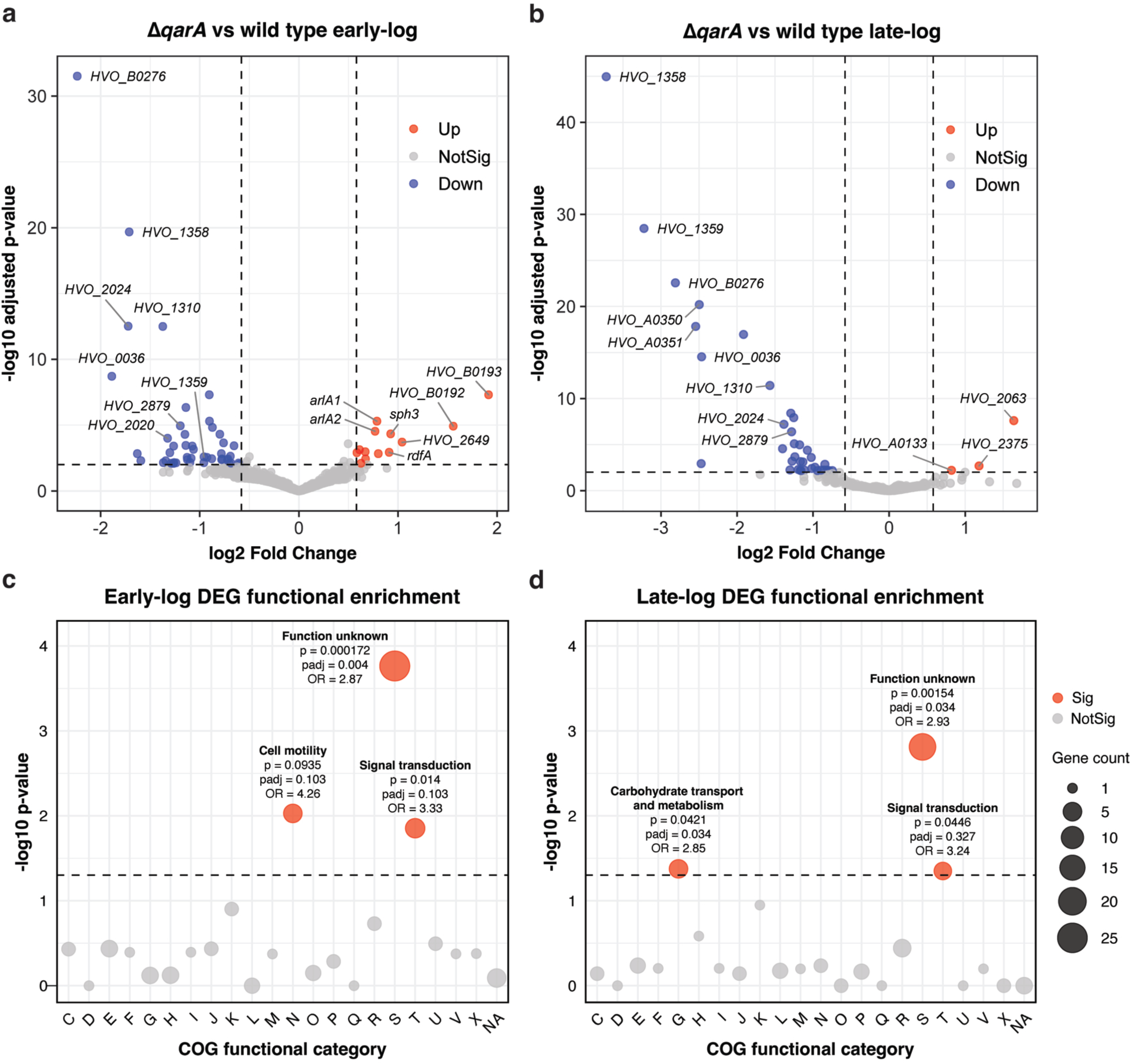
QarA regulates expression of motility and cell-shape determinant genes. Volcano plots depict differential gene expression in *Hfx. volcanii* between Δ*qarA* and wild-type strains during (a) early-log (motile condition) and (b) late-log (non-motile condition) growth. Each point represents a single gene, with genes of interest annotated. Genes with |log₂ fold-change| > 0.58 and adjusted p-value (Benjamini–Hochberg) < 0.01 were considered significantly differentially expressed. *qarA* was excluded from plots to better visualize other DEGs. Bubble plots show significantly enriched COG functional categories among DEGs identified in the Δ*qarA* vs. wild type comparison during early-log (c) and late-log (d) growth. The size of each bubble corresponds to the number of DEGs in the category, and bubble color indicates enrichment significance which was determined by Fisher’s exact test. Categories were considered enriched if they had an unadjusted p-value < 0.05. For significantly enriched categories, the COG function, p-value, adjusted p-value (Benjamini–Hochberg), and odds ratio are reported.

In contrast, 47 genes were downregulated in the Δ*qarA* mutant. A notable finding was the decreased expression of *qarC*. Another gene in this region, *HVO_1359*, encoding a CPxCG zinc-finger protein, was also downregulated. The most strongly downregulated gene overall was *HVO_B0276*, encoding a predicted drug/metabolite transporter (DMT) superfamily protein. Functional enrichment analysis of early-log DEGs revealed, next to motility, an overrepresentation of genes involved in signal transduction (Fig. 7c).

#### Late-log phase

RNA-seq analysis identified 41 DEGs in the Δ*qarA* strain, including 3 upregulated and 38 downregulated genes (Fig. 7b; Table S5). A key observation in late-log was the pronounced downregulation of *qarC* and *HVO_1359*, the latter of which was more strongly down-regulated at increased significance compared to the early-log dataset. *HVO_B0276*, the most strongly downregulated gene in early-log, is also highly downregulated in late-log but drops in relative rank, highlighting the increasing prominence of potential TCS and regulatory components in late-log DFS responses.

The remaining late-log-specific DEGs included upregulation of *HVO_2375* (encoding PstS family transporter), *HVO_2063*, and *HVO_A0133*—encoding a predicted glycine zipper protein with a TAT secretion signal previously reported as highly abundant in both CM-grown and rod-shaped cells (20, 32). Other downregulated late-log-specific genes included *HVO_A0350* (encoding TAT lipoprotein) and *HVO_A0351*. Notably, two mutants with disruptions in *HVO_A0350* were identified in our CM screen (Table 1), supporting a potential role for this region in DFS-dependent phenotypes. Interestingly, our transcriptomics also identified several genes encoding surface-associated PQQ-like β-propeller repeat proteins (*HVO_2606*, *HVO_2607*, *HVO_B0138*, *HVO_B0139*), highlighting potential changes in adhesion, extracellular interactions, or signaling among late-log DEGs. Like in the early-log dataset, functional enrichment analysis found enrichment of genes with roles in signal transduction, but now also carbohydrate metabolism and transport, likely due to surface-associated proteins like the PQQ-like β-propeller repeat proteins (Fig. 7d).

Several genes showed consistent changes across both early- and late-log datasets, suggesting they contribute to a common regulatory program in response to loss of QarA. These included *HVO_1310*, *HVO_0036 and HVO_2879* encoding a probable thioredoxin, probable transmembrane protein, and ornithine cyclodeaminase, respectively. In addition, multiple genes within or near the *HVO_2013–HVO_2020* locus, whose proteins were previously implicated in cell shape determination, were differentially expressed (32). Specifically, *HVO_2020* (encoding a membrane protein; DUF502 family) and *HVO_2024* were downregulated in both datasets, whereas *HVO_2014* and *HVO_2017*, encoding a zinc finger domain protein and a pentapeptide repeat protein, respectively, were identified only in the early-log dataset.

To validate our RNA-seq results, real-time quantitative reverse transcriptase PCR (qRT-PCR) was performed to compare transcript levels between the wild-type and Δ*qarA* strains, focusing on representative genes identified as differentially expressed in the transcriptomic analysis. Consistent with the RNA-seq data, *arlA1* was significantly upregulated in Δ*qarA*, whereas *HVO_B0276* was downregulated (Fig. 8a). Expression of *rdfA* also showed an overall increase in Δ*qarA* (mean relative expression = 1.32); however, variability among biological replicates was high under the tested conditions, and the difference was not statistically significant. Attempts to quantify *qarC* expression were inconclusive due to apparent low transcript abundance (data not shown). Given that Δ*qarC* displays opposite phenotypes to that of Δ*qarA*, and that deletion of *qarA* restores motility in the Δ*qarC* background, we also examined the expression of these same genes in Δ*qarC* to assess potential reciprocal regulation. As expected, *arlA1* and *rdfA* were downregulated in Δ*qarC*, with *arlA1* showing little to no expression, consistent with its non-motile phenotype (Fig. 8b). *HVO_B0276* exhibited a non-significant increase in expression (mean relative expression = 1.4). Strikingly, *qarA* transcript levels were elevated approximately 4 times in Δ*qarC* relative to the wild type, further supporting our hypothesis that QarA is functionally linked to QarB and QarC.

**Figure 8.**
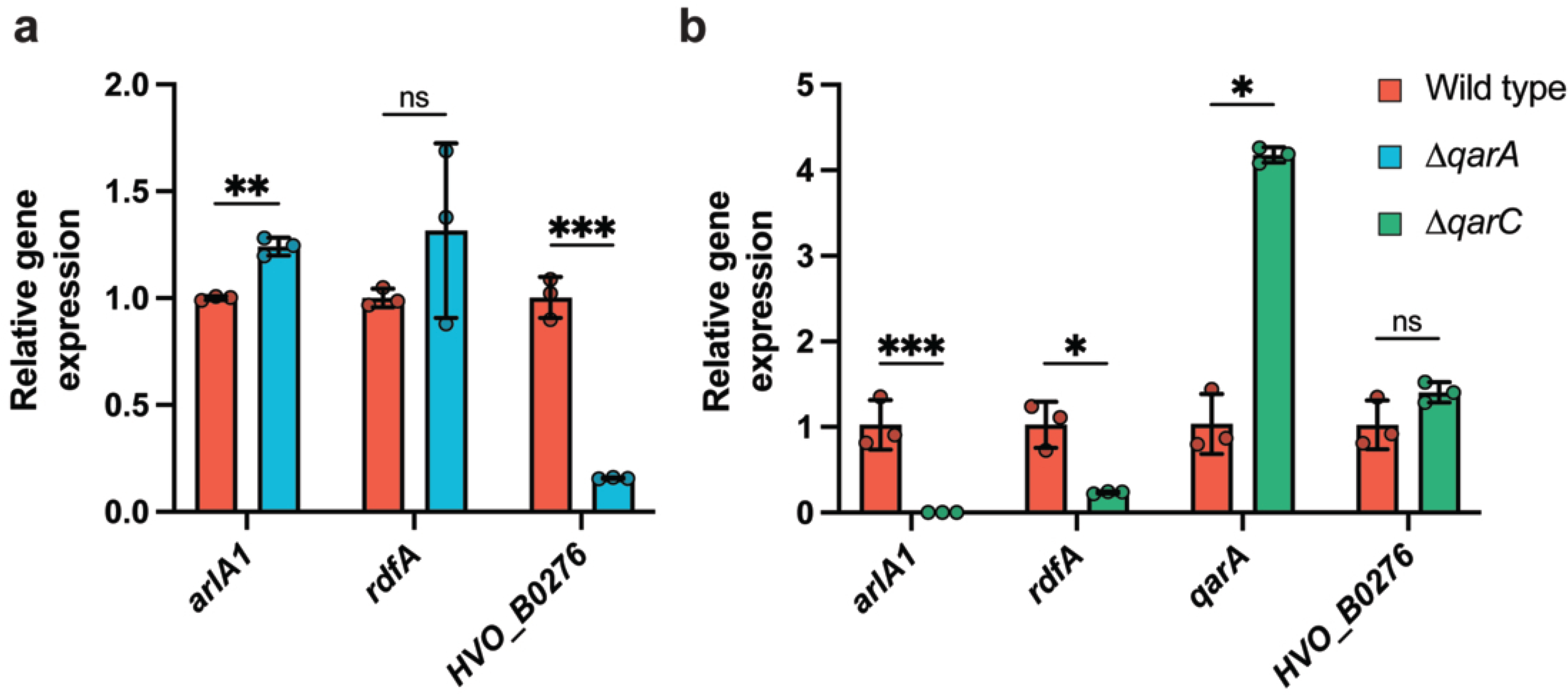
qRT-PCR analysis of Δ*qarA* confirms differential regulation of DEGs identified by RNA-seq while Δ*qarC* displays reciprocal regulation of key DEGs. Reactions were performed using primers for *eif1a*, *arlA1*, *rdfA*, *qarA*, and *HVO_B0276* under the following conditions: (a) wild type vs. Δ*qarA*, and (b) wild type vs. Δ*qarC*. Relative expression levels were calculated using the Pfaffl method, normalized to eif1a, and expressed relative to wild-type controls. Error bars indicate SD of three biological replicates, each with three technical replicates. Statistical significance was determined by log₂-transforming relative expression values and performing unpaired t-tests with Welch’s correction. ns P > 0.05, *P < 0.05, **P < 0.01 ***P < 0.001.

## DISCUSSION

In this study, we developed a genetic screen to identify components of the DFS-mediated QS network in *Haloferax volcanii*. Data obtained from suppressor screens, reverse genetics, microscopy, and transcriptomic analyses, strongly support a three-gene signaling module—*qarABC—*that regulates cell shape and motility in response to DFS. Bioinformatic predictions indicate that *qarA* encodes a multidomain DNA-binding response regulator, *qarB* a cytosolic histidine kinase, and *qarC* a standalone REC-domain protein, all together forming a TCS-like signaling architecture.

Deletion of *qarA* produced a pronounced hypermotile phenotype (Fig. 3a) that does not arise from altered swimming behavior (Fig. 4b, c, d, S3), but instead appears to result from a reduced sensitivity to DFS that regulates the rod-to-disk transition (Fig. 5b). Specifically, Δ*qarA* cells require substantially higher DFS concentrations to fully adopt the disk-shaped, non-motile state, thereby maintaining a larger fraction of rod-shaped, presumably motile cells relative to wild type (Fig. 5e). This reduced sensitivity also explains the enhanced motility observed on soft-agar plates: local DFS accumulation within the halo is less effective at inducing the non-motile state, allowing a greater proportion of cells to remain motile and drive increased migration. Consistent with this interpretation, the DFS concentration present in 25% CM soft agar is likely below the threshold required to fully convert Δ*qarA* cells to disks, permitting continued motility under these conditions.

The role of QarA as a regulator of cell shape is further supported by our finding that in addition to motility-specific genes, expression of key morphological determinants, *sph3* and *rdfA*, is elevated in the absence of *qarA*, suggesting that QarA may act—directly or indirectly—as a repressor of the rod-shaped state in response to DFS levels (Fig. 7a; Table S3). Finally, expression of a QarA variant carrying a substitution at the conserved phosphoacceptor aspartate required for response regulator activation (Fig. S1) failed to complement the Δ*qarA* phenotype in the presence of DFS (Fig. 3d), further supporting the conclusion that QarA functions as a canonical DNA-binding response regulator.

In contrast to the hypermotile Δ*qarA* strain, deletion of *qarB* or *qarC* resulted in non-motile, short rod-shaped cells (Fig. 3a, 4a, 4b) that also transitioned prematurely to disk-shaped cells (Fig. 5b). Like QarA, the conserved phosphoacceptor residues linked to two-component systems in QarB and QarC (Fig. S1) are also required for normal motility responses, supporting their functional role as a histidine kinase and response regulator, respectively (Fig. 3c). Observations that motile suppressors of Δ*qarB* and Δ*qarC* carry secondary mutations in *qarA* (Fig. 6a; Table S1) lead us to perform epistatic analysis of *qarABC*, showing that QarA is indeed epistatic to QarB and QarC, placing QarA as the principal effector of this pathway (Fig. 6b). Moreover, a Δ*qarBC* strain demonstrated identical motility behaviors to its single deletion strains, consistent with a QarB and QarC acting in a shared pathway (Fig. 6b).

### From these data, we propose that QarABC form a DFS-responsive signaling network that governs cell shape transitions (Fig. 9)

Based on their predicted domain architectures, epistasis experiments, and requirement of conserved TCS residues, QarB and QarC likely function as a histidine kinase–response regulator pair, forming the upstream module of this signaling network. Under low DFS conditions, QarB and QarC maintain the motile, rod-shaped state by repressing QarA activity. As DFS accumulates, QarB and QarC become inactivated, relieving repression of QarA. Activated QarA then downregulates rod-associated genes such as *rdfA*, *sph3*, and *arlA1*—which are upregulated in the Δ*qarA* mutant (Fig. 7a, 8a; Table S3)—thereby promoting conversion to the non-motile, disk-shaped state.

**Figure 9.**
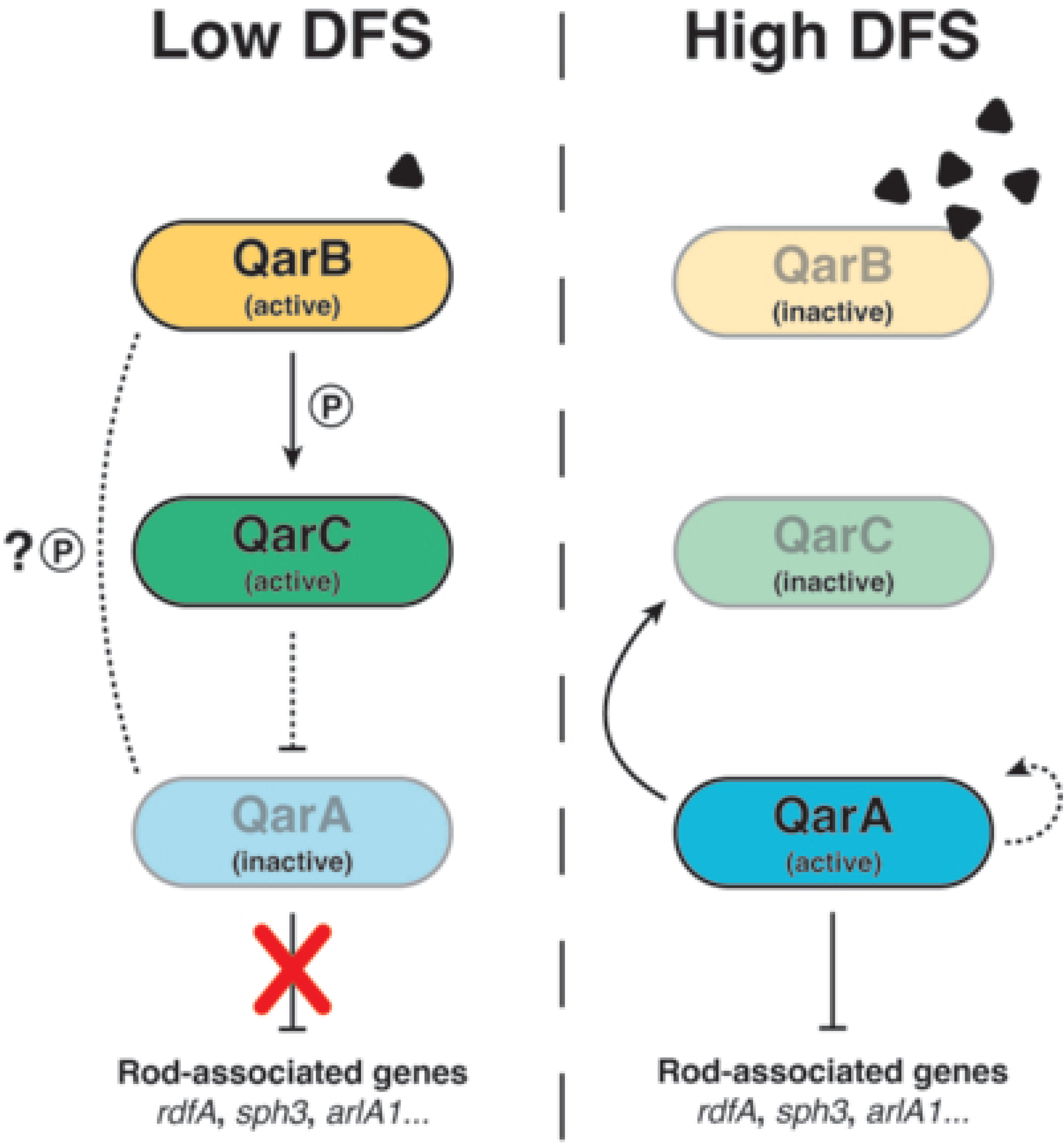
Proposed model of the DFS-responsive QarABC regulatory system. RNA-seq and qRT-PCR data reveal QarA regulatory targets (solid lines from QarA), epistasis experiments and requirement of conserved TCS residues support the upstream QarBC TCS module (solid line between QarB/QarC), while other phenotypic, genetic, and transcriptomic observations support putative interactions (dashed lines).

Consistent with this model, the abundance of QarB was previously observed to decrease under high-DFS conditions (20), suggesting that QarB becomes inactive as DFS levels rise, permitting activation of QarA. Because QarC lacks a canonical output domain (Fig. 2), it is plausible that it transmits its regulatory signal to QarA through direct protein–protein interactions or competition for phosphotransfer from QarB, as QarA also appears to depend on TCS-based phosphorylation for normal motility/cell shape regulation (Fig. 3d)

Beyond this core pathway architecture, we also observe regulatory feedback within the Qar network. Expression of *qarA* is elevated in Δ*qarC* cells (Fig. 7b), which can be explained either by indirect regulation of *qarA* by QarC through other transcriptional regulators, or if QarC is indeed a repressor of QarA, by autoregulation of *qarA* by QarA itself—a mechanism commonly observed among response regulators (37). If QarA indeed activates its own expression, this feedback could amplify its activity once repression by QarB/C is lost. Such a model would explain the reciprocal phenotypes between the Δ*qarB*/Δ*qarC* strains and Δ*qarA*: in both cases, premature QarA activation—driven by loss of upstream inhibition and potential positive feedback—would promote early transition to the non-motile, disk-shaped state even under low DFS conditions. Notably, QarA also promotes expression of *qarC* (Fig. 6a, b), forming a regulatory loop in which QarA potentially activates its own repressor, an arrangement that may serve to fine-tune or stabilize the transition between cell states.

Downstream of QarA, transcriptomic profiling reveals a broader regulatory network linking DFS signaling to motility, morphology, and small-molecule transport. Among the most strongly downregulated genes in Δ*qarA* was *HVO_B0276* (Fig. 7a, b; Table S3, S5), encoding a predicted DMT superfamily transporter previously identified as a regulatory target of the metabolic regulator TbsP, which indirectly influences cell-shape transitions in *Hfx. volcanii* (57). Given that the mechanism of DFS secretion remains unknown, *HVO_B0276* represents a compelling candidate for mediating DFS trafficking.

Two additional genes, *HVO_B0192* and *HVO_B0193*, were strongly upregulated in the *qarA* knockout during early-log phase (Fig. 7a). Both genes are repressed by the cell division regulator CdsR, suggesting that loss of QarA disrupts early shape-related transcriptional programs (58).

Beyond these DFS pathway candidates, we also identified a consistent group of differentially expressed genes across both early- and late-log datasets (Fig. 7a, b), including *HVO_1310, HVO_0036,* and *HVO_2879,* encoding a probable thioredoxin, predicted transmembrane protein, and ornithine cyclodeaminase, respectively, and the *HVO_2013–HVO_2020* region, which encodes multiple shape-related factors in *Hfx. volcanii* (32). Another intriguing candidate is *HVO_1359*, located immediately downstream of *qarABC* and is co-transcribed with *qarC* and *HVO_1360* (Fig. 2)(54, 55). Although *HVO_1359* shows modest differential regulation in early-log phase, it becomes one of the most strongly downregulated genes in late-log phase, suggesting a link to DFS accumulation (Fig. 7a, b). Consistent with this, *HVO_1359* abundance is elevated in cells grown in the presence of high DFS concentrations (20). *HVO_1359* encodes a small CPxCG-type zinc finger protein, part of a conserved but poorly understood family implicated in haloarchaeal stress and signaling regulation (59–61). Notably, *HVO_1359* shares homology and genomic context with the Brz protein of *Hbt. salinarum*, located adjacent to the Bat regulator, with which it is thought to function as a co-regulator of bacterio-opsin transcription (62, 63). This connection suggests that *HVO_1359* could also act as a co-regulator with QarA in DFS signaling. Moreover, zinc finger proteins and AAA-type ATPases, such as *cdc48a* identified in our CM QS bypass screen (Table 1), frequently co-occur with Bat-type proteins in haloarchaeal genomes (63, 64), further suggesting that *HVO_1359* and *cdc48a* may also be components of the DFS regulatory network.

The overall findings described in this work define a DFS-responsive signaling module that integrates environmental and community-derived cues to coordinate collective behavior in *Hfx. volcanii*. The QarABC system represents the first experimentally characterized TCS in Archaea with demonstrated roles in QS-like behavior, exhibiting parallels to both bacterial architectures and archaeal-specific innovations such as standalone REC-domain proteins.

Future studies aimed at dissecting DFS production and sensing, as well as the molecular interactions within the QarABC system, will further clarify the role of QS-like signaling in archaea and its potential conservation across lineages. Although our data support QarABC as a functional two-component–like system, biochemical analyses will be required to define the underlying phosphorylation cascade and establish the directionality of signaling, including direct interactions between QarA, QarB, QarC, and potential other components of this pathway. In addition, while QarA plays a clear role in regulating cell shape and motility responses to DFS, and influences the expression of known cell shape–associated genes, it remains to be determined whether these effects are direct. Establishing direct regulatory targets of QarA will be important to distinguish primary transcriptional control from indirect effects, such as altered cell division or more complex changes in motility behavior (e.g., timing of motility initiation). Finally, while the QarABC system appears to initiate early shape transitions in response to DFS, the ability of Δ*qarA* cells to form disks at high DFS levels indicates parallel pathways governing DFS-dependent cell shape transitions in *Hfx. volcanii* remain to be identified. Together, these directions will illuminate how archaeal communities use small-molecule signaling to regulate collective behaviors, for which the foundations have been laid in the current study.

## METHODS

### Plasmids, primers, and strains

The plasmids, primers, and strains generated through molecular cloning are listed in Table S6. Summarized sequencing results of strains generated through motility screens can be found in Table S1.

### Growth conditions

*Hfx. volcanii* strains were grown at 45 °C in liquid (orbital shaker, 250 rpm, 2.54-cm orbital) or on solid semi-defined Casamino Acids (Hv-Cab) agar (1.5% [wt/vol]) medium (27), supplemented with tryptophan and uracil (50 µg mL⁻¹ each). Uracil was omitted for strains carrying pTA963-based plasmids. Solid plates were incubated in sealable bags to prevent drying. *Escherichia coli* strains were grown at 37 °C in NZCYM medium, either in liquid or solid agar (1.5% [wt/vol]) supplemented with ampicillin (100 µg mL⁻¹). Growth of liquid cultures was monitored via OD₆₀₀ (Thermo Spectronic 20+).

### Preparation of conditioned medium (CM) and CM-agar plates

CM and CM-agar plates were prepared as previously described (20). Briefly, a 1L culture of *Hfx. volcanii* was grown to OD₆₀₀ 1.5 and cells pelleted by centrifugation (Beckman Coulter Avanti J-25I, JA-10 rotor, 8,000 x *g*, 30 min). The resulting supernatant (CM) was then filtered (Millipore Stericup, 0.22 µm PVDF membrane) and stored at −20°C. Soft agar motility plates contained 0.35% [wt/vol] agar of Hv-Cab medium supplemented with CM to a final concentration of 25%.

### Conditioned media motility screens

Colonies of wild type *Hfx. volcanii* were inoculated into 5 mL liquid medium and grown at 45 °C with continuous shaking until mid-log phase (OD₆₀₀ ∼0.5). Assays were conducted using subsurface inoculation or surface inoculation methods, with no comparison being made between the procedures. For isolates beginning with MG (Table S1) cultures were stab-inoculated halfway into the agar with a sterile toothpick, then dragged horizontally within the agar to form a groove. For isolates beginning with SB (Table S1), 20 µL culture was spotted onto CM soft-agar plates and a line streaked across the surface with a sterile toothpick. One or two lines were streaked per plate, followed by incubation at 45 °C for 4–5 days in a plastic box with wet paper towels, changed daily, to prevent drying. Cells from the edge of well-separated outgrowth flares were sampled with sterile toothpicks and streaked on Hv-Cab plates to isolate individual suppressor strains. Colonies were re-stabbed on 25% CM soft-agar plates to confirm the CM-swimming phenotype. Plates were imaged with the lid removed and photographing from above or using a document scanner.

### Whole-genome sequencing of deletion and suppressor strains

Genomic DNA was extracted using the GeneJET Genomic DNA Purification Kit (Thermo Scientific) and quantified with a Qubit 3.0 Fluorometer and BR dsDNA Kit (Invitrogen). Illumina whole-genome sequencing and variant calling were performed by SeqCenter (Pittsburgh, PA, USA). Libraries were prepared using the tagmentation-based Illumina DNA Prep Kit with custom IDT 10 bp unique dual indices and a target insert size of 280 bp. Sequencing was performed on a NovaSeq X Plus, generating 2 × 150 bp paired-end reads. Demultiplexing, quality control, and adapter trimming were performed with bcl-convert (Illumina) (v3.9.3-v4.2.4). Variant calling was carried out with BreSeq (v0.36.1-v0.38.1) using *Hfx. volcanii* DS2 (GCF_000025685.1) as a reference (65). Predicted mutations were compared to the Pohlschröder H53 lab strain (https://doi.org/10.5281/zenodo.19635657) background to identify newly mutated genes. For suppressor strains with no mutations in the, “predicted mutations,” section of the, “index.html,” file provided BreSeq, the “Unassigned missing coverage,” “Unassigned new junction,” sections in the “index.html,” file as well as prections in the “marginal.html” files were manually inspected, prioritizing mutations by their frequency, read quality, as well as DEGs identified in our RNA-seq experiments. Manually predicted mutations were validated by mapping reads to *Hfx. volcanii* DS2 (GCF_000025685.1) using the Geneious mapper (Geneious Prime v.2025.2.2; structural variants, short insertions/deletions of any size, medium/low sensitivity) and confirming the presence of the predicted mutation. Strains with no identified mutations were further screened for large insertions using ISCompare v1.0.5 with default parameters, with none being detected (66).

### Plasmid preparation and transformation

The pTA131 plasmid was used to generate chromosomal deletions (67), and the pTA963 plasmid was used for complementation experiments (68). To prevent restriction of methylated DNA by *Hfx. volcanii*, plasmids were first transformed into *E. coli* NEB 5-alpha, then into homemade *E. coli* DAM⁻ strain DL739, before transformation into *Hfx. volcanii*. Plasmid preparations for both *E. coli* strains were performed using the PureLink Quick Plasmid Miniprep Kit (Invitrogen). *Hfx. volcanii* transformations were carried out using the polyethylene glycol method (69), and *E. coli* transformations by heat shock (70).

### Generation of chromosomal deletions in *Hfx. Volcanii*

Chromosomal deletions were generated using the pop-in/pop-out method following transformation with pTA131-based plasmids, as previously described (67). Constructs to generate knockout strains (pJC1, pJC2, pJC3) were generated by amplifying ∼750 nt flanking each gene via PCR, followed by overlap PCR (primer sets 3–4, 7–8, 11–12). The fused upstream and downstream fragments were cloned into pTA131 at XbaI and XhoI sites. Following transformation and selection, deletions were confirmed by colony PCR using primers flanking and internal to the deleted gene (primer sets 5–6, 9–10, 13–14). Deletion strains were further validated by whole-genome sequencing and variant calling to confirm complete gene deletion and assess secondary genome alterations. The double deletion strain Δ*qarBC* (Δ*HVO_1356*Δ*HVO_1358*) was generated by transforming pJC2 (pTA131::Δ*HVO_1356*) into the Δ*qarC* (Δ*HVO_1358*) background as previously described.

### Construction of complementation plasmids

The HVO_1356-StrepII expression plasmid (pJC4) was generated by PCR amplification of *HVO_1356* from *Hfx. volcanii* genomic DNA using primer set 15, followed by cloning into pTA963. The corresponding complementation plasmid, HVO_1356-StrepII-Myc (pJC6), was constructed by introducing a Myc tag into pJC4 via site-directed mutagenesis (Q5 Site-Directed Mutagenesis Kit, NEB) using primer set 16. Similarly, HVO_1357-StrepII (pJC5) was generated by PCR amplification with primer set 17 and cloning into pTA963. The complementation plasmid HVO_1357-StrepII-FLAG (pJC7) was then created by introducing a FLAG tag into pJC5 using site-directed mutagenesis with primer set 18. Phosphoablative mutants of HVO_1356 and HVO_1357 were generated by substituting alanine at positions D67 and H347, respectively. These mutations were introduced into pJC6 and pJC7 via site-directed mutagenesis as described above, using primer sets 19 and 20, yielding plasmids pJC10 and pJC11. The complementation plasmid HA-StrepII-HVO_1358 (pJC9) and its corresponding D58A phosphoablative mutant (pJC12) were synthesized by Twist Bioscience (San Francisco, CA, USA). All complementation constructs contain a 1× StrepII tag (WSHPQFEK), together with either a 1× Myc (QarB/HVO_1356; EQKLISEEDL), 1× FLAG (QarA/HVO_1357; DYKDDDDK), or 1× HA (QarC/HVO_1358; YPYDVPDYA) tag for future biochemical assays. The tags are C-terminal for QarA/QarB and N-terminal for QarC. The StrepII tag is separated from the protein by a GGP linker and from the detection tag by a GGGGS linker.

### Growth curves

Colonies of each strain were inoculated into 5 mL liquid medium and grown at 45 °C with continuous shaking until mid-log phase (OD₆₀₀ 0.3–0.5). Cultures were diluted to OD₆₀₀ 0.01 in 200 µL fresh medium per well of a non-treated, flat-bottom 96-well plate (Corning). Growth was measured using a BioTek Epoch 2 microplate reader with Gen6 software (Agilent v1.04). Four biological replicates were used per strain. To minimize evaporation, empty wells were filled with 200 µL water, and a 2 °C gradient was applied to the 45 °C incubator setting. Readings were taken every 30 min at 600 nm, with continuous double-orbital shaking (335 rpm, 4 mm) between measurements, for a total of 72 h.

### Determination of viable cell counts

Cultures for each strain were grown to mid-log phase (OD₆₀₀ 0.3–0.5). Cultures were diluted to OD600 0.3 and serially ten-fold diluted. 30µL of 10^−3^ to 10^−5^ dilutions were spread onto Hv-Cab solid agar plates and incubated for 3-5 days. Colonies were manually counted and reported as log10 CFU/mL of the original culture.

### Motility assays and motility halo quantification

Motility was assessed on 0.35% [wt/vol] agar Hv-Cab medium. Colonies were inoculated into 5 mL liquid medium and grown at 45 °C with continuous shaking until mid-log phase (OD₆₀₀ 0.3–0.5). Cultures were diluted to the lowest OD₆₀₀ among the strains tested and stab-inoculated into the agar with a toothpick. Plates were incubated upright at 45 °C in a plastic box with wet paper towels, changed daily, to maintain humidity. After two days, plates were removed and imaged following at least one day at room temperature to allow halos to darken. Images were acquired using a document scanner, with lids removed. Halo areas were quantified as previously described (57) using FIJI (71): the scale was calibrated to plate diameter, halos were segmented with the threshold tool, and areas measured using the Analyze Particles function.

### Cell shape microscopy

*Hfx. volcanii* strains were inoculated from single colonies into 5 mL Hv-Cab liquid medium; for CM experiments, 50 µL CM (1% [vol/vol]) was added. Cultures were grown to early-log (OD₆₀₀ 0.045–0.050), mid-log (OD₆₀₀ 0.30), or stationary phase (72 h, OD₆₀₀ ∼1.9–3.0). For early- and mid-log cultures, 1 mL aliquots were centrifuged at 4,900 × g for 6 min and pellets resuspended in their supernatant (∼5 µL for early-log, ∼50 µL for mid-log). For imaging, 1.5 µL of resuspended pellet or stationary culture was placed on glass slides (Fisher Scientific) and covered with No. 1 18 × 18 mm coverslips (Globe Scientific Inc.), gently pressed to maintain a single cell layer. Images were acquired on a Leica DMi8 inverted microscope with DFC9000 GT camera using Leica Application Suite X (v3.6.0.20104). DIC and brightfield images were captured with a 100× Plan Apochromat objective (NA 1.4, WD 0.09 mm). Brightfield images for quantification were background-subtracted in FIJI (ImageJ v2.16.0/1.54p)(71) and analyzed in CellProfiler (v4.2.8)(72). The FIJI macro and CellProfiler pipeline used are available at https://doi.org/10.5281/zenodo.19635657.

### CM titration microscopy

*Hfx. volcanii* strains were inoculated from single colonies into 5 mL Hv-Cab liquid medium containing 0.2, 0.6, 1, 2, or 4µL/mL CM. Cultures were grown to early-log phase (OD₆₀₀ 0.045–0.050) and imaged and analyzed using FIJI (ImageJ v2.16.0/1.54p)(71) and CellProfiler (v4.2.8)(72) as described above. Fraction of disk-shaped cells was calculated by finding the ratio of cells with the CellProfiler eccentricity values below 0.9.

### Motility chamber construction

Motility chambers were assembled using 24 × 50 mm (Fisher Scientific) and 18 × 18 mm (Corning) No. 1 coverslips. Coverslips were cleaned in a saturated potassium hydroxide/ethanol bath for 30 min, rinsed with deionized water, and immersed in 1% solution of polyvinylpyrrolidone K30 containing 0.01% Tween-20 for 1 h. Coverslips were removed slowly, air-dried vertically, and baked at 120 °C for 1 h. Chambers were constructed by placing two double layers of parafilm (∼5 × 25 mm) ∼8 mm apart on a prepared 24 × 50 mm coverslip on a 95 °C hotplate, followed by covering them with a prepared 18 × 18 mm coverslip to form an ∼8 × 18 mm chamber with a height of ∼200 µm.

### Live-cell tracking and analysis

Cultures were grown to early-log phase (OD₆₀₀ 0.045–0.050) and diluted 1:5 (wild type, Δ*qarA*) or 1:2 (Δ*qarB*, Δ*qarC*) in pre-heated (45°C) motility buffer (Hv-Cab medium with 1% [wt/vol] BSA). Motility chambers were partially filled (∼2/3) from one end with pre-heated motility buffer to further passivate glass surfaces, then fully filled with diluted cells. Chamber ends were sealed with liquid VALAP wax and imaged immediately. Swimming behavior was recorded on a Nikon TI2-E microscope with a 20× CFI60 Plan Apochromat Lambda objective (NA 0.75, WD 1.0 mm) under brightfield illumination. Videos were captured at 5 fps for 5 min using a Hamamatsu Orca-Fusion Gen-III sCMOS camera using NIS-Elements (6.10.01). Cells were tracked near the bottom coverslip. Videos were processed in FIJI, and trajectories were automatically analyzed using a custom macro with TrackMate plugins (71, 73). For determination of non-motile vs motile cells, 1*µm*/s was used as a threshold based on observations of non-motile objects in the motility chambers.

### RNA extraction and purification

Liquid cultures (5 mL) of each strain were grown at 45 °C with shaking to OD₆₀₀ 0.05 (early-log) or 0.9 (late-log). Early-log (4–5 mL) and late-log (0.5 mL) cultures were harvested and pelleted at 4,900 × g for 6 min. RNA was extracted using the Qiagen RNeasy Plus Micro Kit, with lysates homogenized via QIAshredder columns. For late-log RNA-seq and qRT-PCR samples, DNase digestion was performed using the RNase-free DNase kit (Qiagen) followed by RNeasy MinElute Cleanup (Qiagen). RNA concentration was measured with Qubit RNA Broad Range Assay Kits (Invitrogen), and integrity assessed using RNA ScreenTape on a 4200 Tapestation (Agilent). All RNA-seq and qRT-PCR samples had RINe values above 8.9 and 7.0, respectively.

### RNA sequencing and data analysis

Cell pellets were collected and RNA extracted as described previously. For early-log and late-log samples, four and three biological replicates were collected, respectively. Purified RNA was sent to SeqCenter LLC (Pittsburgh, PA, USA) for library preparation and sequencing. RNA was DNase-treated (Invitrogen, RNase-free) prior to library preparation using Illumina Stranded Total RNA Prep Ligation with Ribo-Zero Plus kit and 10 bp unique dual indices. Additional rRNA depletion was performed with *Hfx. volcanii*-specific probes from Rados *et al*. (74). Sequencing was performed on a NovaSeq X Plus with 2 × 150 bp paired-end reads. Demultiplexing, quality control, and adapter trimming were done with bcl-convert (v4.2.4). Paired-end RNA-seq reads were quality-checked, rRNA/tRNA reads were removed, and the remaining reads were aligned to the *Hfx. volcanii* DS2 reference genome (GCF_000025685.1) using minimap2 (75), with alignments processed using samtools (76). Gene-level counts were obtained with featureCounts from the Rsubread R package (77). Differential expression between Δ*qarA* and wild type was analyzed using DESeq2 (78); genes with <10 total counts were excluded. Log₂ fold changes were estimated with the Wald test and shrinkage-adjusted using apeglm (79). Gene annotations, including product descriptions, arCOG assignments, and COG functional classes, were obtained from RefSeq and locus tag reference tables. Genes with adjusted p-values (Benjamini–Hochberg) <0.01 and |log₂FC| > 0.58 were considered differentially expressed. PCA and volcano plots were generated with ggplot2 and plotly to visualize sample clustering and differential expression. Functional enrichment of significant DEGs was assessed by comparing the distribution of gene function categories in each DEG set to the genome-wide background using Fisher’s exact tests. The pipeline used to perform RNA-seq analyses can be found at https://doi.org/10.5281/zenodo.19635657

### qPCR analysis

Three biological replicates of Δ*qarA* and Δ*qarC* were collected as previously described for early-log RNA samples, along with two sets of three biological replicates of wild-type controls per comparison. cDNA was synthesized from 1 µg RNA using the ProtoScript II First Strand cDNA Synthesis Kit (New England BioLabs) with random primers; parallel negative controls without reverse transcriptase (-RT) were prepared.

Primers for *Hfx. volcanii* genes *eif1a, arlA1, rdfA, HVO_1357, HVO_1358,* and *HVO_B0276* were designed using Primer3 and synthesized as RxnReady™ primers. qRT-PCR reactions (20 µL) contained 2× iTaq Universal SYBR Green Supermix (Bio-Rad), 5% DMSO, 300 nM primers, and 8 µL of 1:125 diluted cDNA. No-template (NTC) and -RT controls were included; reactions were considered acceptable if ΔCt ≤10 between sample and control. Primer efficiencies determined via standard curves ranged from 85.5–99.5%. Each reaction included three biological replicates, each with three technical replicates, run on a QuantStudio 3 Real-Time PCR System (Applied Biosystems).

Thermal cycling conditions were: 94 °C for 2 min, then 40 cycles of 95 °C for 15 s and 60 °C for 30 s (*eif1a*, *HVO_1357*, *HVO_B0276*) or 58 °C for 30 s (*arlA1*, *rdfA, HVO_1358)*. SYBR Green fluorescence was recorded at each cycle’s end, and the default melt curve program following the qPCR run verified primer specificity. Relative expression was calculated in Excel using the Pfaffl method (80), normalizing to *eif1a*, and compared to wild-type controls to determine differential expression.

### Multiple sequence alignments

Protein sequences of experimentally characterized archaeal histidine kinases and response regulators (25, 41–43), along with the CheY response regulator and EnvZ histidine kinase from *Escherichia coli*, the BAT protein from *Hbt. salinarum*, and QarABC proteins, were retrieved from UniProt (81). Sequences were manually curated to retain only regions corresponding to the REC (IPR001789) or HIS_KIN (IPR005467) domain profiles as defined in InterPro (48). The resulting sequences were grouped by protein class (response regulators or histidine kinases) and aligned using Clustal Omega with default parameters (82). Alignments were subsequently visualized in Jalview (83).

## DATA AVAILABILITY

The raw RNA-seq data generated in this study have been deposited in the NCBI Sequence Read Archive (SRA) under BioProject accession number PRJNA1347779. Additional materials are available at https://doi.org/10.5281/zenodo.19635657, including: whole-genome sequencing and variant-calling data for mutant, knockout, and background strains; images, scripts, and processed data for cell-shape analyses; images and data for soft-agar motility assays; live-cell tracking data; pipelines and key outputs from RNA-seq analyses; raw and processed qPCR data; and multiple sequence alignments.

## ACKNOWLEDGEMENTS

We acknowledge Kiara Escudero, Shalaya Brown, and Sophia Sun for their contributions to phenotypic assays and CM screens. We also thank Dr. Alexandre Bisson for his guidance with RNA-seq analysis and for providing rDNA-depletion probes. We gratefully acknowledge Dr. Friedhelm Pfeiffer, Dr. Fevzi Daldal, Dr. Yirui Hong, and Everlyne Mutua for their insightful feedback on the manuscript. This work was supported by the National Science Foundation (UPenn NSF-MBC2222076) and the University of Pennsylvania Research Fund. J.C. was supported by the National Science Foundation Graduate Research Fellowship. P.C. was supported by a National Institutes of Health Training Grant (UPenn T32 GM007229) as well as a National Institutes of Health Ruth L. Kirschstein National Research Service Award (1F31AI181536-01). M.G. was supported by a Merck/PennFERBS fellowship. A.M. acknowledges funding from the Charles E. Kaufman Foundation (Early Investigator Research Award KA2022-129523; New Initiative Research Award KA2024-144001) and the National Science Foundation (UPenn MRSEC DMR-2309043).

