## Supplemental Figures for "A Two-Component Regulatory System Mediates Quorum Sensing–Dependent Morphology and Motility Transitions in the Archaeon *Haloferax volcanii*"

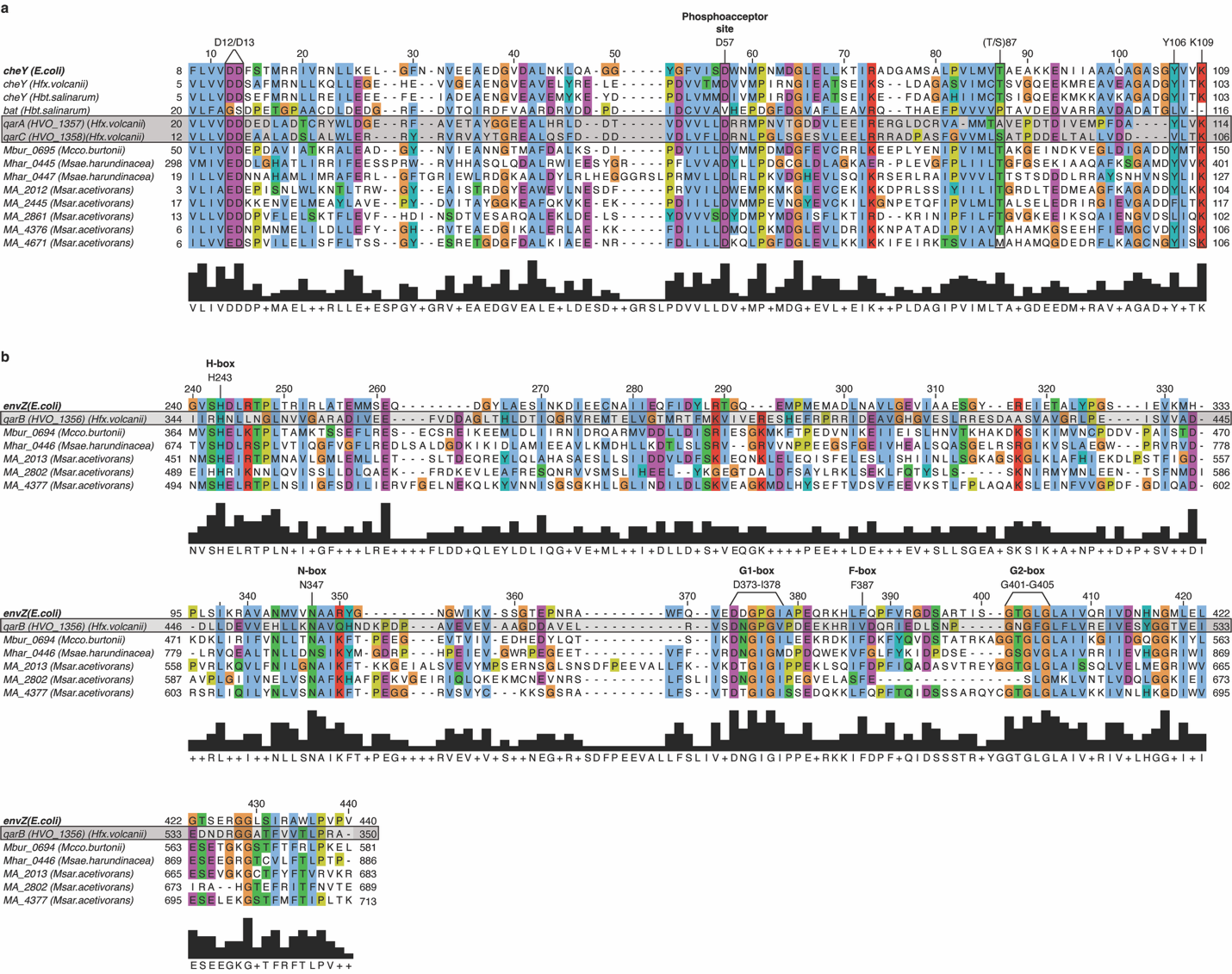

**Figure S1. Multiple sequence alignment of *Hfx. volcanii* histidine kinase and response regulator proteins (QarABC) with representative archaeal and Escherichia coli sequences.** Alignments were visualized in Jalview, with shading based on the default ClustalX coloring scheme. A conservation histogram and consensus sequence are shown below each alignment. In panel (a), key catalytic residues of *E. coli* CheY (D12–D13, D57, T/S87, Y106, K109; UniProt: P0AE67) are indicated above the alignment, with residue numbering referenced to CheY. In panel (b), conserved motifs of *E. coli* EnvZ (H-box, N, G1, F, and G2 boxes; UniProt: P0AEJ4) are highlighted above the alignment, with residue numbering referenced to EnvZ. CheY D57 corresponds to D67 in QarA and D58 in QarC, and EnvZ H243 corresponds to H347 in QarB.

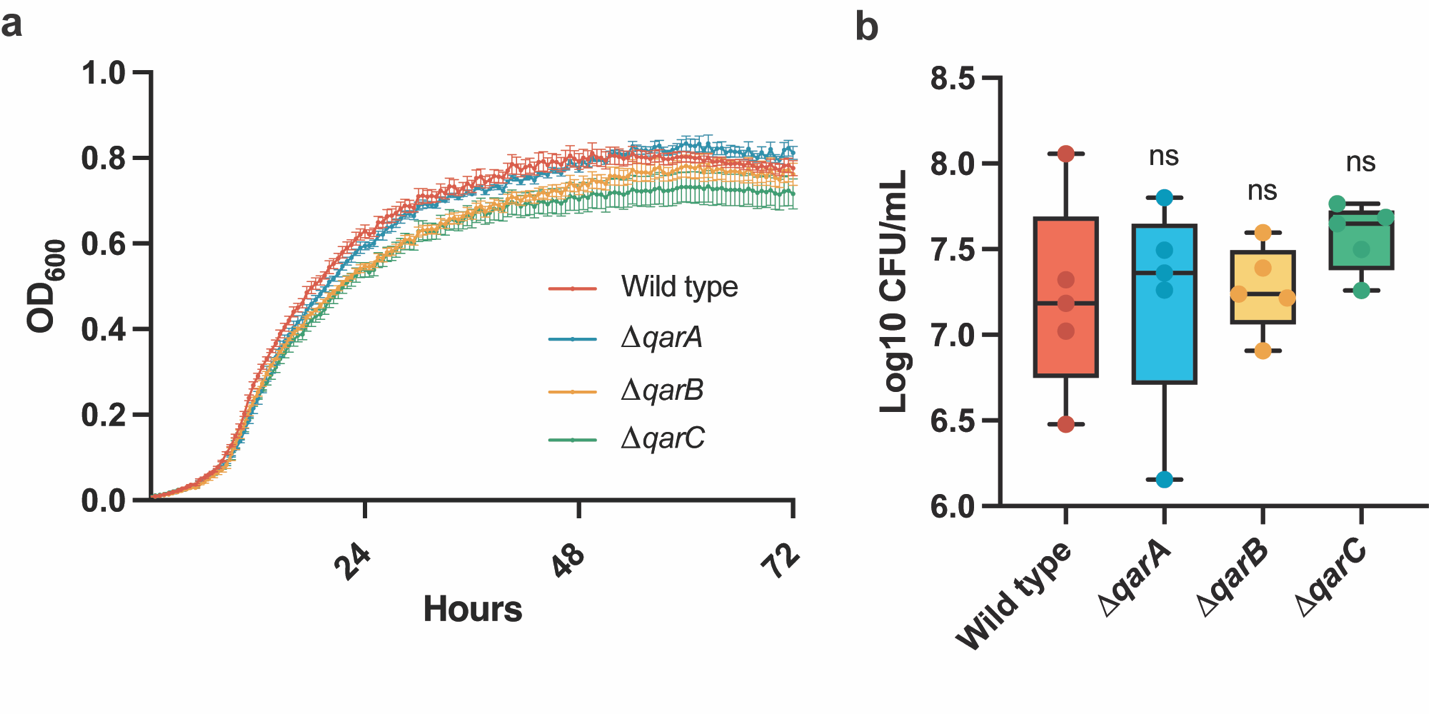

**Figure S2. Growth and viability of QarABC mutants.** (a) Growth curves of wild-type and QarABC mutants cultured in liquid medium at 45 °C for 72h. Data represent the mean and SD of four biological replicates. (b) Viable cell counts (log₁₀ CFU/mL) of mid-log-phase (OD_600_ 0.3) cultures determined by serial dilution and plating on Hv-Cab agar. Error bars represent SD of five biological replicates from five independent experiments.

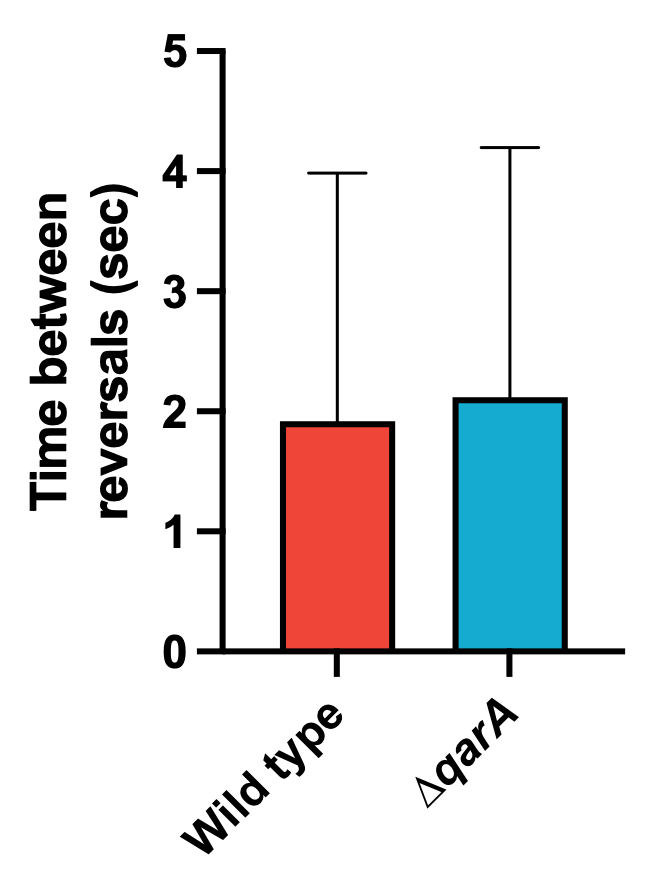

**Fig. S3. ∆*qarA* appears to have a similar reversal frequency to wild type.** Cells were grown to early-log phase (OD₆₀₀ = 0.045–0.050) and diluted 1:5. Diluted cultures were loaded into pre-prepared motility chambers (see Methods) and imaged immediately under brightfield illumination. Swimming behavior was recorded at 5 fps for 5 min, and trajectories were extracted in FIJI and analyzed using a custom TrackMate-based macro. This resulted in >2000 trajectories for each strain from two independent experiments. Reversal frequency was determined by the average time between two subsequent >90° turns for each tracked cell over its entire track. Error bars represent SEM.
